# Deletion of CEP164 in mouse photoreceptors post-ciliogenesis interrupts ciliary intraflagellar transport (IFT)

**DOI:** 10.1101/2022.03.21.485080

**Authors:** Michelle Reed, Ken-Ichi Takemaru, Guoxin Ying, Jeanne M. Frederick, Wolfgang Baehr

## Abstract

Centrosomal protein 164 (CEP164) is located at the edge of distal appendages in primary cilia and is necessary for basal body (BB) docking to the apical membrane. To investigate the function of CEP164 before and after BB docking in photoreceptors, we deleted CEP164 during retina embryonic development (Six3Cre), in postnatal rod photoreceptors (iCre75) and in mature retina using tamoxifen induction. BBs dock to the cell cortex during postnatal day 6 (P6) and extend a connecting cilium (CC) and an axoneme. P6 retina-specific knockouts (^ret^*Cep164*^-/-^) are unable to dock BBs, thereby preventing formation of a CC or outer segments (OS). In rod-specific knockouts (^rod^*Cep164*^-/-^), Cre expression starts after P9 and CC/OS form. P16 ^rod^*Cep164*^-/-^ rods have nearly normal OS lengths, and maintain OS attachment through P21 despite loss of CEP164. Intraflagellar transport components (IFT88, IFT57 and IFT140) were reduced at P16 ^rod^*Cep164*^-/-^ BBs and CC tips and nearly absent at P21, indicating impaired intraflagellar transport. Nascent OS discs, labeled with a fluorescent dye on P14 and P18 and harvested on P19, showed continued ^rod^*Cep164*^-/-^ disc morphogenesis but absence of P14 discs mid-distally, indicating OS instability. Tamoxifen induction with PROM1^ETCre^;*Cep164*^F/F^ (^tam^Cep164^-/-^) adult mice affected maintenance of both rod and cone OS. The results suggest that CEP164 is key towards recruitment and stabilization of IFT-B particles at BB/CC. Impairment of IFT may be the main driver of ciliary malfunction observed with hypomorphic CEP164 mutations.

**Author summary:** Centrosomal protein of 164 kDa (CEP164) is indispensable for docking of basal bodies to apical membranes and formation of primary cilia. Homozygous truncations of CEP164 are associated with severe ciliopathy whereas missense mutations generate milder forms of NPHP. Recessive mutations of CEP164 are associated with nephronophthisis (NPHP), Meckel-Gruber (MKS) and Bardet-Biedl syndromes (BBS) in which primary cilia fail to form, or do not function correctly. We found that deletion of CEP164 in mouse photoreceptors before docking of the basal body (BB) to the apical inner segment membrane prevents elaboration of connecting cilia (CC, equivalent to transition zones) and outer segments (OSs, modified primary cilia). We also generated mouse models in which deletion of CEP164 occurs after OS are established and observed that the BB/CC and axoneme are initially stable in the absence of CEP164. Loss of CEP164, however, diminished recruitment of IFT-B proteins to the BB/CC affecting intraflagellar transport (IFT), which is required for maintenance of cilia and OSs. We propose that, in ciliopathies caused by missense mutations (e.g., X1460WextX57 associated with LCA) impairment of IFT provides a mechanism for ciliary dysfunction and disease.

## Introduction

CEP164 (NPHP15) is a ubiquitously expressed centrosomal protein [1] participating in microtubule organization, cell cycle progression and basal body (BB) docking. CEP164 localizes to the BB distal appendages (DAPs) which protrude from the BB distal end as nine-bladed pinwheel-like structures that enable BB docking and ciliogenesis (**Fig. S1A**) [2, 3]. Additional key DAP proteins are SCLT1, CEP83, CEP89, FBF1 and C2CD3 at the distal BB center, proteins containing multiple coiled-coil domains [2, 4]. STED microscopy demonstrated that CEP164 localizes to the outermost of several concentric rings surrounding the BB distal end [5, 6]. CEP164 recruits TTBK2 (tau tubulin kinase 2) to the BB, an interaction based on the CEP164 WW domain with a TTBK2 proline-rich domain [7] (**Fig. S1B**). CEP164-TTBK2 interaction facilitates TTBK2-mediated phosphorylation of both CEP83 [8, 9] and MPHOSP9 (M-Phase Phosphoprotein 9, MPP9) at the distal end of the mother centriole [10]. MPHOSP9 and the associated CP110-CEP97 complex that caps the distal ends of mother centrioles are removed subsequently to promote ciliary axoneme growth [10-12].

Recessive mutations of CEP164 (e.g., Q11P, R93W, Q525X, R576X, R579X) **Fig. S2A**) are associated with nephronophthisis (NPHP), Meckel-Gruber syndrome (MKS) and renal-retinal ciliopathies [13]. Homozygous truncations of CEP164 are associated with severe ciliopathy whereas missense mutations generate milder forms of NPHP [1, 13]. Homozygous CEP164(Q11P) and compound R93W with Q525X are associated with NPHP, while homozygous CEP164(R93W) also has been identified in a patient with BBS [13, 14]. Both mutations are outside the WW domain located between residues 56 and 89 [12], but structural analysis revealed that Q11P and R93W mutations compromise TTBK2 complex formation [7]. CEP164(*X1460WextX57*), identified in a child with LCA [13], causes a read-through of the stop-codon X1460, adding 57 foreign amino acid residues to the CEP164 C terminus. Biallelic pathogenic variants in CEP164 are causative of Oral-facial-digital syndromes (OFDVI) [15].

Knockdown of CEP164 in zebrafish resulted in syndromic ciliopathy with ventral body axis curvature, cell death, abnormal heart looping, pronephric tubule cysts, hydrocephalus, and retinal dysplasia [13, 16]. Homozygous *Cep164* mouse knockouts were embryonically lethal [17]. In a conditional knockout mouse model that lacks CEP164 in multiciliated tissues and testes, a profound loss of airway, ependymal, and oviduct multicilia resulted, and the mutant mouse developed hydrocephalus and male infertility [17]. Using tracheal cell cultures of this mouse model, CEP164 was shown to regulate small vesicle recruitment, ciliary vesicle formation and BB docking [17].

Photoreceptor OSs are considered modified primary cilia, dedicated to light reception and phototransduction [18-21]. As CEP164 forms a complex with multiple ciliary proteins [22], multiple functions may be revealed depending on time of depletion. We generated CEP164 conditional knockouts in retina with three different *Cre* drivers that produce CEP164 truncation during retina embryonic development (^ret^*Cep164*^-/-^) pre-ciliogenesis, in rod photoreceptors (^rod^*Cep164*^-/-^) post-ciliogenesis, and in adult retina using tamoxifen induction (^tam^*Cep164*^-/-^). ^ret^*Cep164*^-/-^ mice were unable to elaborate photoreceptor CC and OS, whereas ^rod^*Cep164*^-/-^ mice do form photoreceptor CC and OS as *Cre* expression starts post-ciliogenesis. Depletion of CEP164 post-ciliogenesis results in profound loss of IFT-B particles, leading to impairment of intraflagellar transport (IFT) and OS degeneration.

## Results

### Generation of retina-specific *Cep164* knockout mice

The murine *Cep164* gene (30 exons, one noncoding exon 1) produces multiple splice variants lacking internal exons. The main variant encodes a protein of 1333 amino acids (150 kDa) carrying an N-terminal WW domain and three coiled-coil domains (**Fig. S2A**). To generate a retina-specific knockout mouse at various stages of development, *Cep164*^F/F^ mice [17] (**Fig. S2B**) were bred with transgenic Six3Cre mice (^ret^*Cep164*^-/-^) [23], iCre75 mice (^rod^*Cep164*^-/-^) [24], or Prom1(C-L) (Prom1-ETCre) (^tam^*Cep164*^-/-^) mice [25] to produce null alleles (**Fig. S2C**). Six3 (*sine oculis*-related homeobox 3) is a transcription factor expressed in retina, RPE and brain during embryonic development [23]. *Cre* in iCre75 is under control of the rhodopsin promoter that expresses in the second postnatal week, allowing CC and OS to develop. Tamoxifen induction allows depletion of CEP164 at any time during development including in the adult mouse. LoxP sites placed in introns 3 and 4 specify deletion of exon 4 (**Fig. S2B**), truncating CEP164 after exon 3 (after residue Pro 64) as exon 5 is out-of-frame. Truncation after exon 3 is predicted to eliminate all splice variants, including the WW domain interacting with TTBK2. Genotyping with tail and retina DNA as templates and primers flanking exon 4 verified knockout of the *Cep164* gene (**Fig. S3**). Immunohistochemistry with anti-CEP164 antibody showed that CEP164 is a photoreceptor inner segment (IS) protein and accumulates at the photoreceptor BB, identified by EGFP-CETN2 (**Fig. S3G**, upper panel) in P6 wildtype (WT) retina. In contrast, CEP164 was undetectable at BBs of ^*ret*^*Cep164*^*-/-*^ central retina (**Fig. S3G**, lower panel).

### ^ret^*Cep164*^-/-^ photoreceptors lack connecting cilia and outer segments

Six3*Cre* expression begins at embryonic day 9 (E9) and the floxed gene of interest is deleted subsequently. In P6 *Cep164*^*F/+*^;Egfp-Cetn2 controls, BBs docked to the cell cortex as evidenced by presence of CC distal to BBs (**Fig. 1A**). CEP164 appears firmly attached to BBs (**Fig. 1A’**, arrows) and short CC are formed (**Fig. 1A’**, arrowheads). Few ^ret^*Cep164*^-/-^ BBs carry CEP164 and CC formation declines in central ^ret^*Cep164*^-/-^ photoreceptors (**Fig. 1B**), presumably because BBs are unable to dock to the IS apical membrane. Similarly, in multiciliated Cep164^F/F^;FoxJ1-Cre MTEC cells (mouse tracheal epithelial cells), up to 83% of BBs were undocked [17].

**Figure 1.**
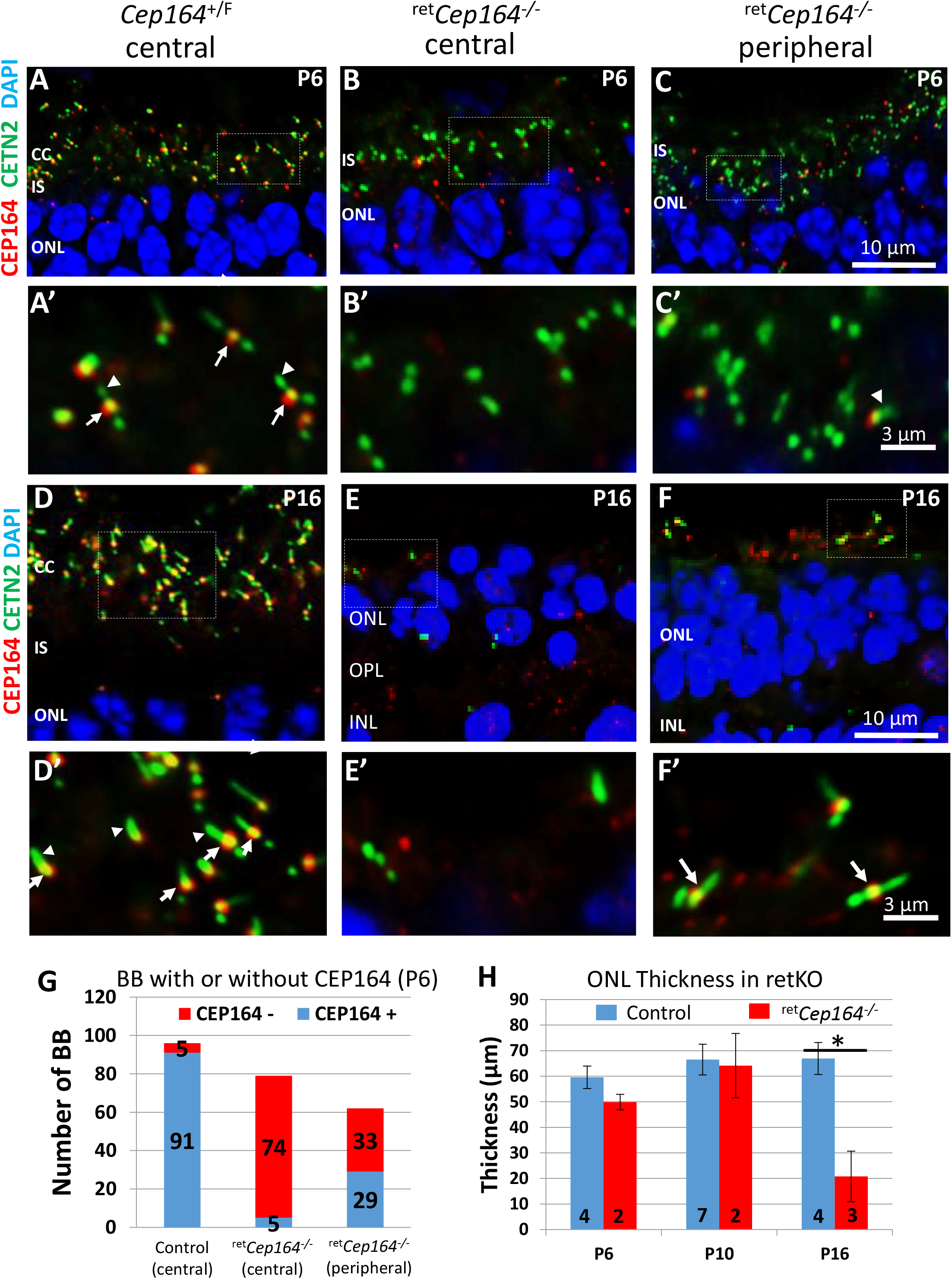
Rapid postnatal ^ret^*Cep164*^*-*/-^ photoreceptor degeneration. **A-C**, P6 *Cep164*^+/F^ (A), ^ret^*Cep164*^*-/-*^ central (B) and peripheral (C) cryosections on Egfp-Cetn2^+^ background probed with anti-CEP164 antibody (red). In ^ret^*Cep164*^*-*/-^ cryosections, CEP164-BB association is much reduced, but not in retina periphery where degeneration is delayed. **A’-C’**, enlargements of A-C as indicated. CEP164 associates with the BB (arrows) in control sections (A’), is absent in the knockout (B’), but detectable at some BB in the retina periphery (C). Arrowheads identify CC. **D-F**, P16 cryosections probed with anti-CEP164 antibody. Note reduced thickness of mutant ONL. **D’–F’**, enlargements as indicated in D-F. At P16, control CC (D) are normal length and CEP164 is firmly associated with the BB (arrows). In central ^ret^*Cep164*^*-*/-^ sections (E), ONL is reduced to 1-3 nuclear rows, BBs and CCs have disintegrated. In the peripheral retina (F), few BB structures survive (arrow). **G**, number of BBs in P6 ^ret^*Cep164*^*-/-*^ and control photoreceptors. In P6 ^ret^*Cep164*^*-*/-^ retina cryosections, CEP164 association with BB is much reduced compared to CEP164^+/F^ control (n =5 and 91 BBs, respectively) and CC are not formed. **H**, ONL thickness at P6, P10 and P16 in central ^ret^*Cep164*^*-/-*^ (n = 2-7). Note significant decrease at P16.

Six3Cre expression is delayed in peripheral ^ret^*Cep164*^*-/-*^ retina (**Fig. 1C**), and consequently some photoreceptors develop short CC (fewer than 10 in >50 BBs) (**Fig. 1C’**, arrowhead). Mother centrioles with bound CEP164 (**Fig. 1D’**, arrows) extend fully developed CC in P16 control photoreceptors (**Fig. 1D’**, arrowheads), whereas in central knockout retina, BBs disintegrate and photoreceptors degenerate (**Fig. 1E, E’**). Few BBs with CC survive in the retina periphery (**Fig. 1F** and **F’**, arrow). Quantitative evaluation reveals that 91 of 96 control BBs, but only 5 of 79 knockout BBs, were labeled by antibody directed against CEP164 in P6 central retina (**Fig. 1G**). Absence of 100% labeling in the control is attributed to the imaging plane since CC formation was robust. Thickness of ^ret^*Cep164*^-/-^ ONL appears to be stable at P6 and P10, but declines by P16 (**Fig. 1H**).

### ^ret^*Cep164*^*-/-*^ photoreceptor morphology

Plastic sections of P7 control and ^*ret*^*Cep164*^*-/-*^ retinas show that the ONL and IS are comparable (**Fig. 2A, B**). Ultrastructure of a representative P7 ^*ret*^*Cep164*^*+/-*^ photoreceptor reveals a BB docked to the cell cortex with a CC (**Fig. 2C**). By contrast, ^*ret*^*Cep164*^*-/-*^ BBs were missing distal appendages, failed to dock to the plasma membrane and aborted ciliogenesis (**Fig. 2D**).

**Figure 2.**
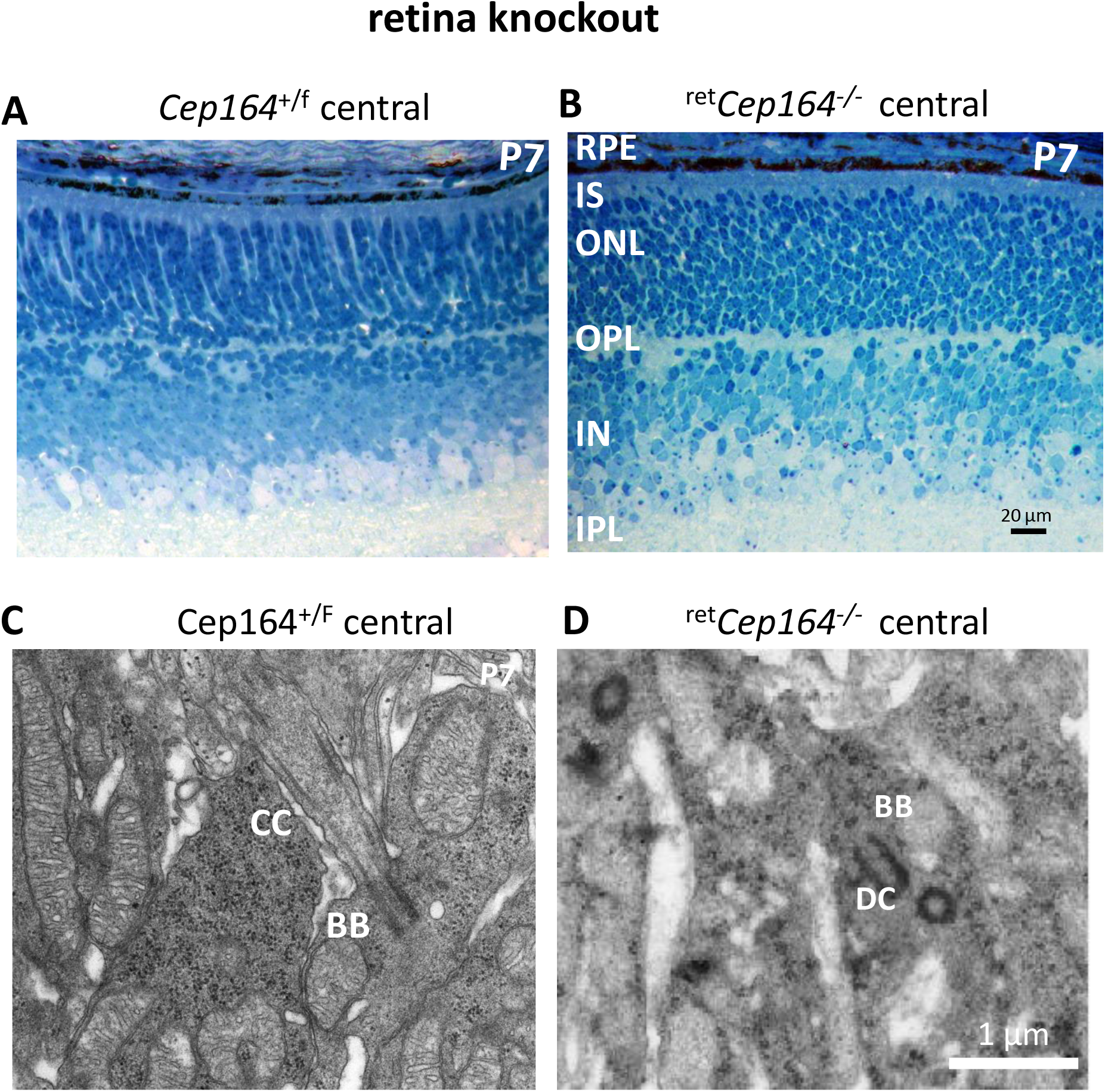
P7 retina histology and photoreceptor ultrastructure. **A, B**, plastic sections of control (A) and ^ret^*Cep164*^-/-^ retina (B) show normal histology. **C, D**, Ultrastructure reveals docked BB, CC formed in control (C). In the knockout, BB/DC is not associated with apical membranes and CC is not formed (D).

### ^ret^*Cep164*^*-/-*^ photoreceptor function at P16 and P25

As Six3Cre expression generates a central-to-peripheral gradient of gene product depletion [26], antibodies directed against PDE6 and ARL13B were used to assay for OS elaboration in central versus peripheral retina. PDE6 is a key phototransduction enzyme [27] and ARL13B is a small GTPase functioning as a guanine nucleotide exchange factor (GEF) of ARL3 [28]. In cryosections of *Cep164*^+/F^;Egfp-Cetn2 control retina, PDE6 (MOE antibody) and ARL13B localize in the OS as expected (**Fig. 3A, D**). In central ^ret^*Cep164*^*-/-*^ sections, PDE6 (**Fig. 3B**) and ARL13b (**Fig. 3E**) mislocalize to the IS as OSs fail to form. In peripheral retina (**Fig. 3C, F**), PDE6 and ARL13B localize to rudimentary OS. OS formation in the retina periphery encouraged functional analysis by pan-retina electroetinography (ERG). Scotopic a-wave (rod function) and photopic b-wave amplitudes (cone function) with flashes of 1.4 log cd s·m^-2^ are almost extinguished at P16 and P25 (**Fig. 3G, H**). High flash intensities under scotopic conditions elicit weak a-wave responses (**Fig. 3I**) and under photopic conditions an even stronger response (**Fig. 3J**) consistent with minor OS formation and photoreceptor survival at the retina periphery.

**Figure 3.**
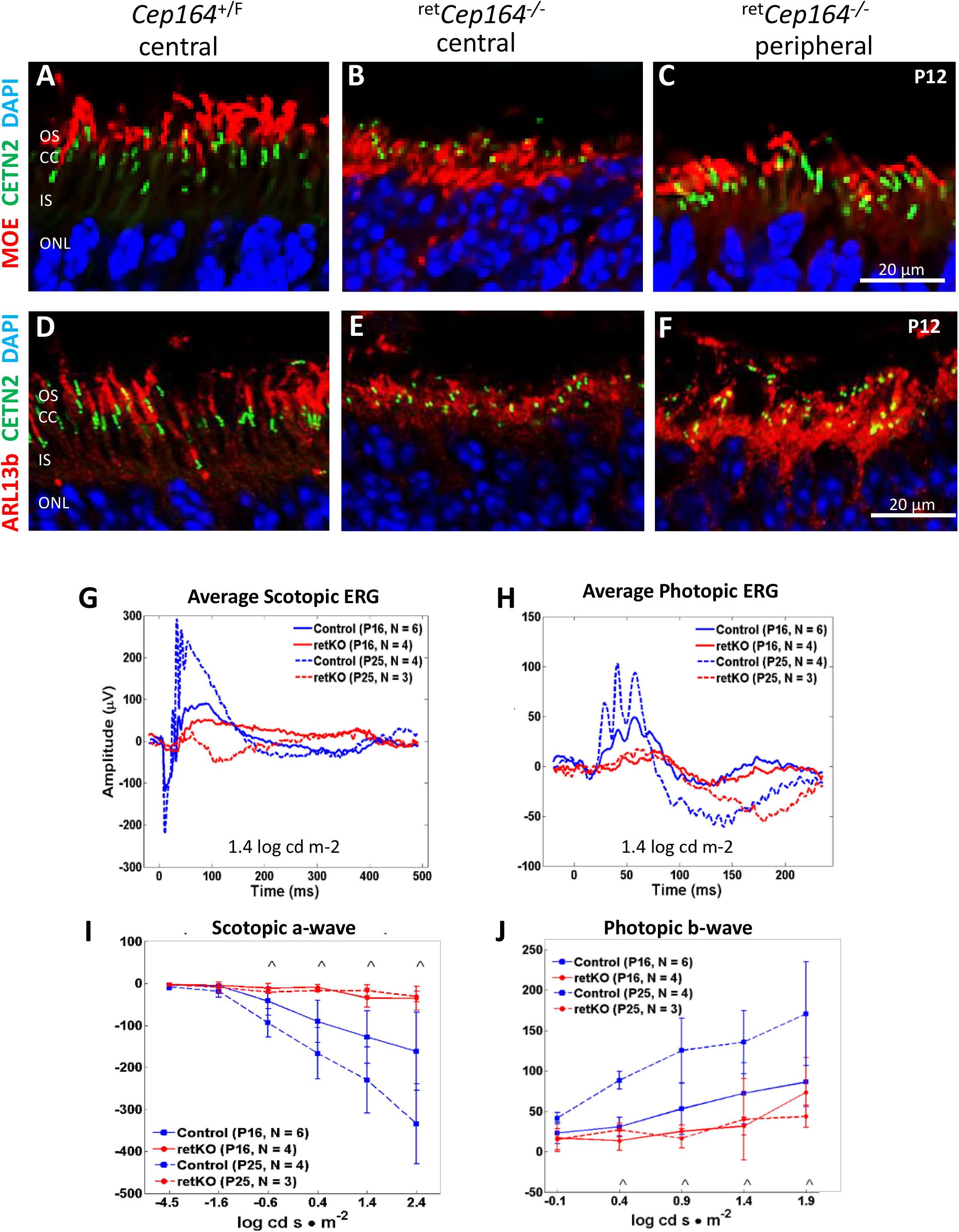
Central ^ret^*Cep164*^-/-^ *photoreceptors* do not form OS. **A-C**, P12 central *Cep164*^+/F^ (A), central ^ret^*Cep164*^*-/-*^ (B) and peripheral ^ret^*Cep164*^*-/-*^ retina cryosections(C) probed with MOE (anti-PDE6) (red). PDE6 is an OS protein participating in phototransduction. In central ^ret^*Cep164*^*-/-*^ sections (B), OS are not formed and PDE6 mislocalizes to the IS. In retina periphery (C), degeneration is delayed and some OS are present. **D-F**, cryosections as in A-C probed with anti-ARL13B, a small GTPase present in the OS. In controls (D), ARL13B correctly localizes to OS, whereas it mislocalizes to photoreceptor IS of central and peripheral ^ret^*Cep164*^*-/-*^ retina. **G-J**, Electroretinography (ERG) at P16, P25. Average scotopic ERG (G) and photopic ERG (H) in control (blue) and retina knockouts (red) at 1.4 log cd s m^-2^. Scotopic a-wave (G) and photopic b-wave (H) amplitudes (μV) as a function of light intensity (log cd s m^-2^). Scotopic ERGs show diminished responses by P16 and statistically different responses (^, p < .05) at P25. Photopic ERGs show diminished b-wave responses by P16 and statistically different responses (^, p<.05) at P25. Minimal scotopic a- and b-waves reveal surviving photoreceptors of retina periphery.

### Cep164 rod knockout

Previous reports of CEP164 dealt with its roles in early steps of ciliogenesis such as preciliary vesicle recruitment and BB docking, but its role in ciliary maintenance is unknown. To determine whether CEP164 is required once CC are established, we examined deficits in a rod-specific knockout of CEP164 driven by iCre75 under control of the rhodopsin promoter (**Fig. S2**). One advantage of a rod-specific knockout is that *Cre* expression does not occur before P10, thus permitting ciliogenesis and OS formation. We asked whether CEP164 is necessary for continued rod viability and, if so, how fast rods would degenerate after CEP164 deletion.

Genotyping of ^rod^*Cep164*^*+/-*^ control and *Cep164*^F/F^;iCre75 (abbreviated ^rod^*Cep164*^*-/-*^) retina DNA confirmed that *Cep164* exon 3 excision was complete at P16 (**Fig. S3H**, lane 5). Plastic sections of P16 ^rod^*Cep164*^*-/-*^ photoreceptors demonstrated that IS/OS length and ONL thickness are comparable to heterozygous control at P16 (**Fig. 4A**), but reduced at P21 (**Fig. 4B**).

**Figure 4.**
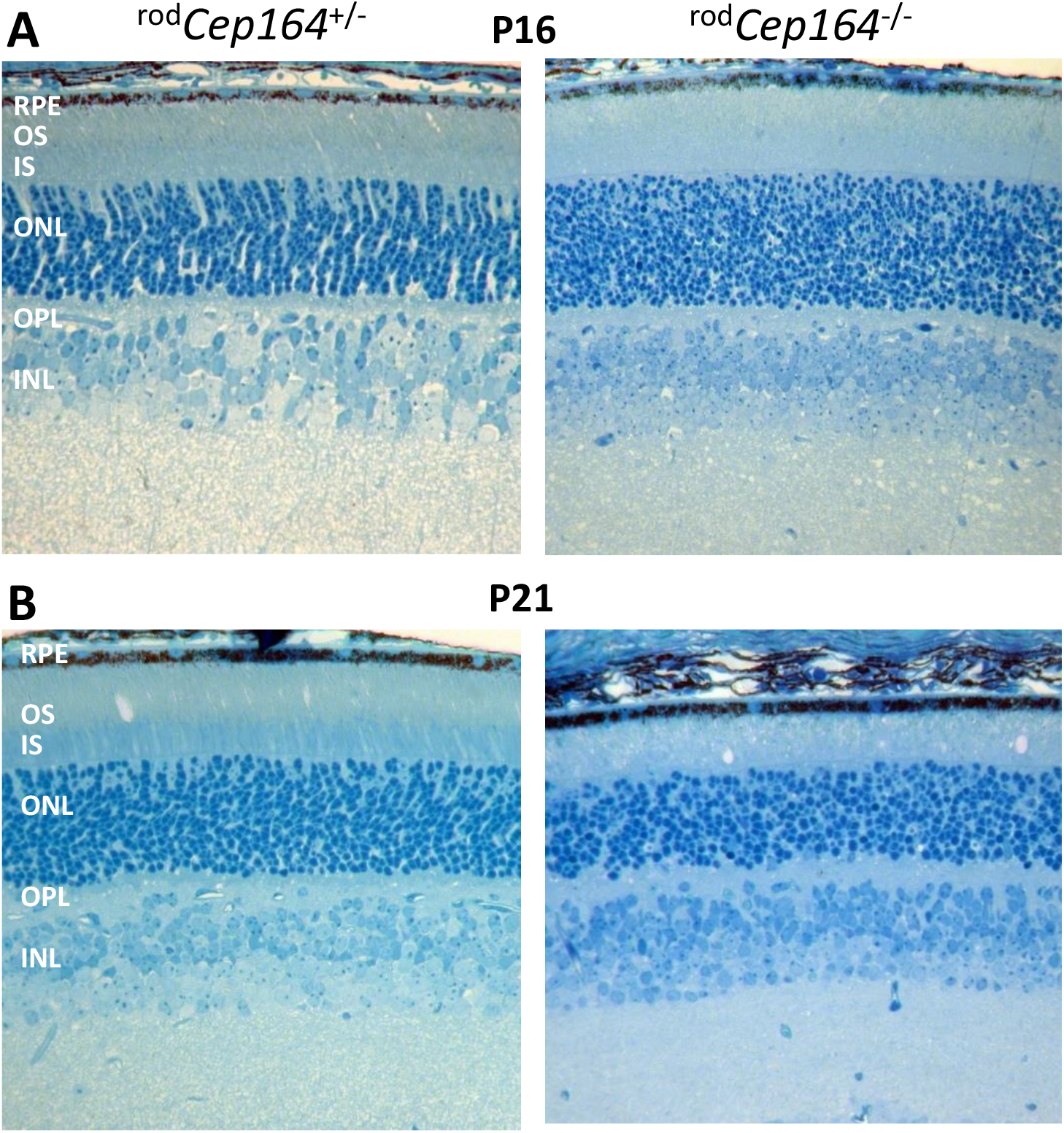
^rod^*Cep164*^-/-^ retina histology. **A, B**, retina morphology at P16 (A) and P21 (B). Note OS are still full-length at P16, but reduced by P21.

Immunohistochemistry at P15, P21 and P30 showed that CEP164 was firmly attached to the BB distal appendages in *Cep164*^+/F^;Egfp-Cetn2 cryosections (**Fig. 5**, row **A** and **A’**). At P15, ^rod^*Cep164*^*-/-*^ BBs have lost most CEP164 and at P21, CEP164 was undetectable at BBs (**Fig. 5**, row **B, B’**). The P30 knockout ONL revealed one layer of nuclei; a single BB/CC structure is likely that of a surviving cone (**Fig. 5B’**, P30; **Fig.7H**).

**Figure 5.**
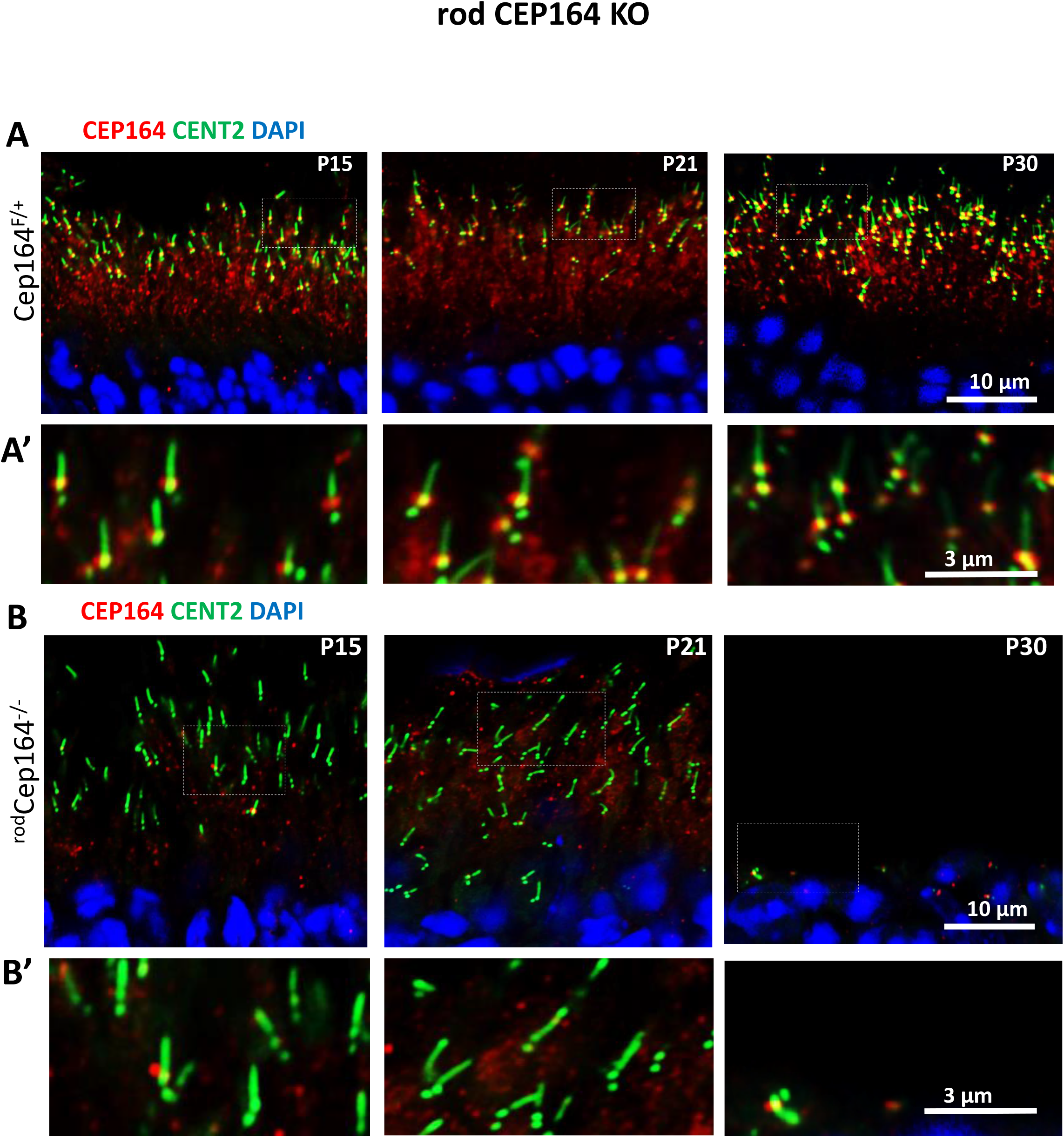
Cep164 rod knockout and BB/CC stability. Row **A**, *Cep164*^+/F^ ;iCre75 (^rod^*Cep164*^+/F^) retina cryosections on Egfp-Cetn2^+^ background probed with CEP164 antibody (red) at P15, P21 and P30. Row **A’**, enlargements as indicated by dashed squares in A. Row **B**, ^rod^*Cep164*^-/-^ cryosections probed with CEP164 antibody. Row **B’**, enlargements as indicated by dashed squares in B. Note rapid rod degeneration from P15-P30, but BB and CC appear perfectly normal at P15 and P21 in the absence of CEP164. Scale bar, 10 μm in row A and B, 3 μm in row A’, B’.

### ^rod^*Cep164*^*-/-*^ outer segment instability

Depletion of CEP164 after P16 raises the question of BB-membrane attachment and OS stability. We again used anti-PDE6 and anti-ARL13B antibodies to identify OS proteins by immunohistochemistry (**Fig. 6A**). At P16, ^rod^*Cep164*^*-/-*^ OSs appeared to be stable in the absence of CEP164 (**Fig. 6B**). However, by P21, OS became unstable and the ONL thickness was reduced to one-third (**Fig. 6B, D**). By P30, ^rod^*Cep164*^*-/-*^ OSs and ONL are undetectable. At P16, ARL13B is barely detectable in the ^rod^*Cep164*^*-/-*^ OS and is absent in the mutant OS at P21 (**Fig. 6D**), suggesting that CEP164 is necessary for ARL13B to reach the WT OS. While the ^rod^*Cep164*^*-/-*^ ONL was not significantly reduced at P12 and P16 relative to control ONLs (**Fig. 6E**), the P16 ^ret^*Cep164*^*-/-*^ ONL thickness at P16 was less than half compared to control consistent with distinct onset of Cre-based recombination (**Fig. 1H**).

**Figure 6.**
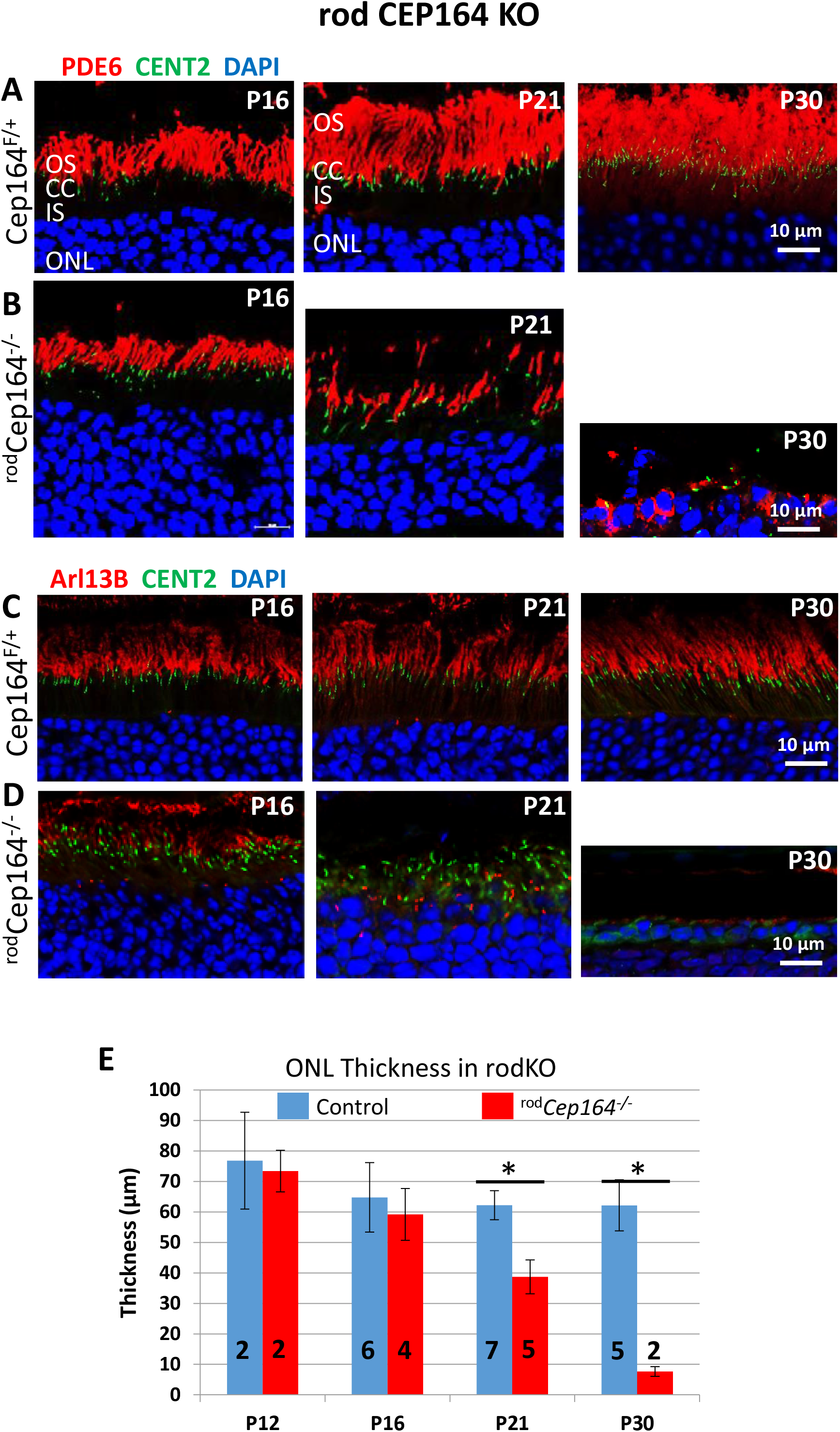
^rod^*Cep164*^-/-^ OS degenerate after P16. Rows **A, B**, *Cep164*^+/F^ (A) and *Cep164*^*F/F*^;iCre75 (^rod^*Cep164*^-/-^) retina cryosections (B) probed with PDE6 antibody (red) (A, B) at P16, P21 and P30. All sections are on Egfp-Cetn2 background. Note disintegrating ^rod^*Cep164*^-/-^ OS in B at P21 where CC (green) is still congruent with the OS (red). Rows **C, D**, central *Cep164*^+/F^ (C) and central ^rod^*Cep164*^*-/-*^ (D) retina cryosections probed with anti-ARL13B (red) at P16, P21 and P30. Note ARL13B, known to interact with CEP164, does not reach the OS in the absence of CEP164. **E**, ONL thickness at P12, P16, P21 and P30 in central retina of ^rod^*Cep164*^-/-^ (n = 2-7). ONL thickness is indistinguishable at P12 and P16, but decreases rapidly after P16.

### ^rod^*Cep164*^*-/-*^ photoreceptor function

Scotopic and photopic pan-retina ERGs were performed at a constant 1.4 log cd s/m^2^ intensity and the traces averaged (N=4-14) (**Fig. 7A, D**). In scotopic ERG, control rod a-waves continue to increase from P15 (120 μV) to P30 (300 μV) while ^rod^*Cep164*^-/-^ rod a-waves hover at 50 μV at P16 and P21 (**Fig. 7A, B**). At P30, the ^rod^*Cep164*^-/-^ rod a-wave is nearly extinguished consistent with absence of rods (**Fig. 6B**). The photopic ^rod^*Cep164*^*-/-*^ cone b-wave is identical to control at P15, reduced at P21 and extinguished at P30, indicating cone degeneration. This is consistent with previous findings where deletion of rod-specific genes resulted in secondary loss of cones due to diminished rod-derived cone viability factor (RdCVF) [29]. Scotopic ERG as a function of light intensity showed diminished rod a- and b-wave amplitudes as early as P15 beginning at 1.4 log cd s/m^2^ (**Fig. 7C**) suggesting rod degeneration has begun, although not statistically significant at that age. Photopic cone b-wave amplitudes of control and rod knockout mice were identical at P15 but those of knockouts were nearly extinguished by P30 (**Fig. 7E, F**). Assuming that cone bipolar cells are unaffected in ^rod^*Cep164*^-/-^ retinas, the severely attenuated photopic cone b-wave derives from loss of cone function. Cone degeneration at P30 was verified using anti-ML-opsin antibody (**Fig. 7G, H**). P21 ^rod^*Cep164*^-/-^ cones are relatively stable, but disintegrate by P30.

**Figure 7.**
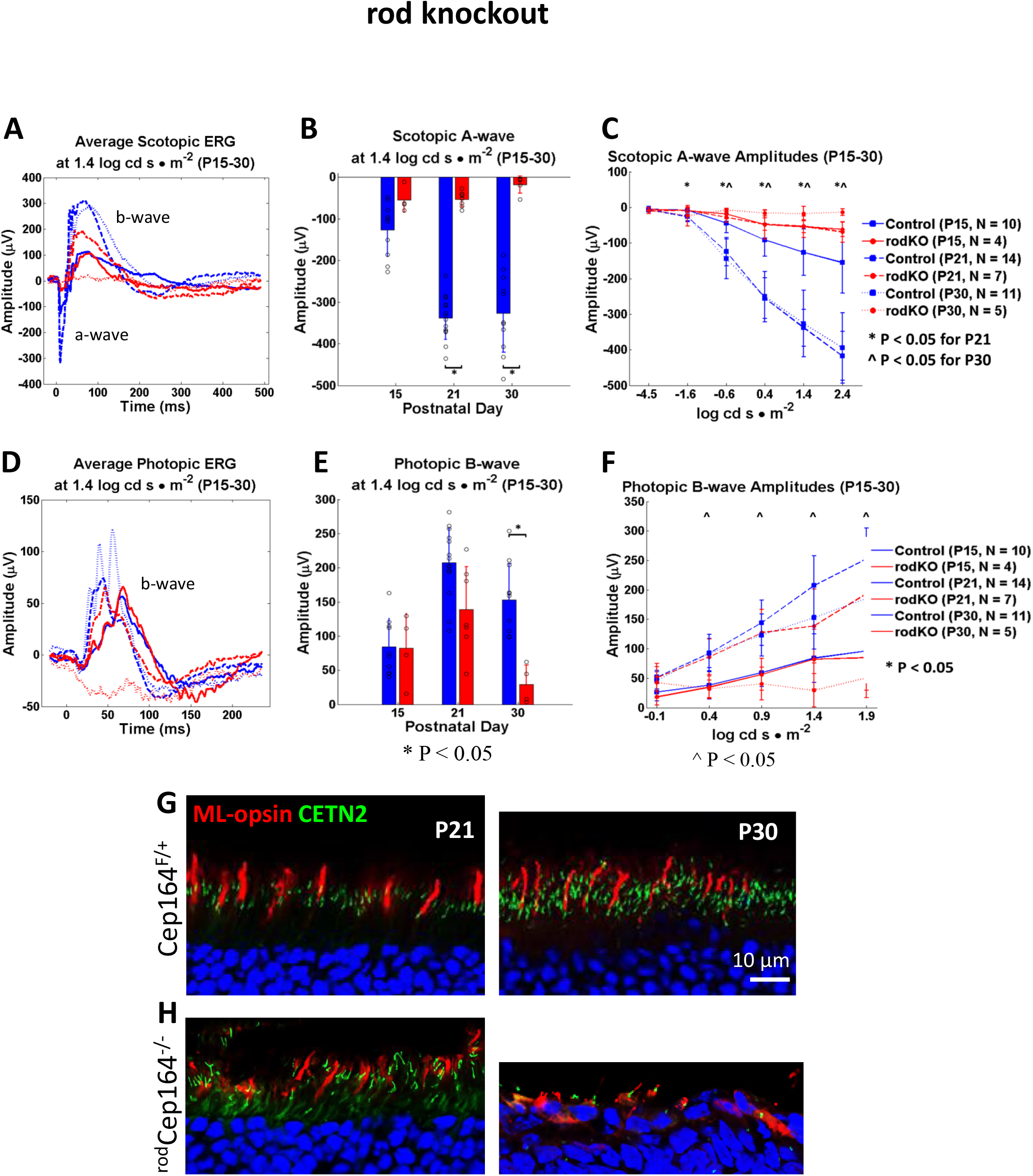
^rod^*Cep164*^-/-^ scotopic and photopic ERG. **A**, control (blue) and ^rod^*Cep164*^-/-^ mice (red) were dark-adapted overnight and exposed under scotopic conditions to flashes of 1.4 log cd s/m^2^ at P16, P21 and P30. Note knockout a- and b- wave amplitudes were diminished at all ages. **B**, statistical evaluation of amplitudes of control and knockout CEP164 scotopic ERG a-waves at P15, P21 and P30 as measured in A. **C, F**, scotopic a-wave (C) and photopic b-wave amplitudes (F) at P16, P21 and P30 as a function of light intensity. Knockout scotopic a- and b-waves are diminished, but never extinguished. **D, D**, control (blue) and ^rod^*Cep164*^-/-^ mice (red) were exposed under photopic conditions to flashes of 1.4 log cd s/m^2^. Note, ^rod^*Cep164*^-/-^ photopic b-waves were absent at P30. **E**, statistical evaluation of photopic b-wave amplitudes at 1.4 log cd s/m^2^ as measured in B. **G, H**, immunohistochemistry with anti-ML-opsin of control (G) and ^rod^*Cep164*^-/-^ central cryosections (H) at P21 and P30. Cones on the ^rod^*Cep164*^-/-^ background are stable until P21 but are degenerated at P30.

### Loss of IFT88, IFT57 and IFT140 at BB and CC

In primary cilia (RPE1 cells), CEP164 recruits NPHP1 and IFT88 to the mother centriole and, upon deletion of CEP164, the levels of BB-associated IFT88 strongly decreased [30]. In contrast to primary cilia, CEP164 is dispensable for the recruitment of IFT components to multicilia [17]. We explored whether IFT88, IFT57 and IFT140 levels vary in P16 ^rod^*Cep164*^-/-^ retinas before onset of degeneration, and at P21 when OSs disintegrate and ONL thickness is reduced (**Fig. 6**). P16 and P21 control ^rod^*Cep164*^+/-^ photoreceptors reveal the familiar pattern of IFT proteins positioned at the BB and CC distal tips (**Figs. 8A-E**, upper panels) [31-33]. By contrast, P16 ^rod^*Cep164*^-/-^ photoreceptors reveal reduced levels of IFT88 and IFT57 at BB and CC (**Figs. 8A-B**, lower panels), and IFT protein levels are further attenuated in P21 knockouts (**Figs. 8C-F**, lower panels). SPATA7 (spermatogenesis associated protein 7) [34] and CEP290 [35, 36] are unaffected by deletion of CEP164 (**Fig. 9A, B**) suggesting normal CC elaboration. NPHP1 remains unaffected by post-ciliogenesis deletion of CEP164 (**Fig. 9C**). FBF1 remains at the BB in the ^rod^*Cep164*^*-/-*^, suggesting intact DAPs (**Fig. 9D**). These results suggest that CEP164 recruits IFT88, IFT140 and IFT57 to photoreceptor BB/CC, as observed in primary cilia [30]. IFT88 and IFT57 protein levels are reduced in the ^rod^Cep164^-/-^, negatively affecting IFT and OS axoneme stability as evidenced by shortening of ^rod^*Cep164*^*-/-*^ rod OS at P16 and P21 (**Fig. 6B, D**) and absence of P14 discs in the dye injection experiment (see below, **Fig. 10**).

**Figure 8.**
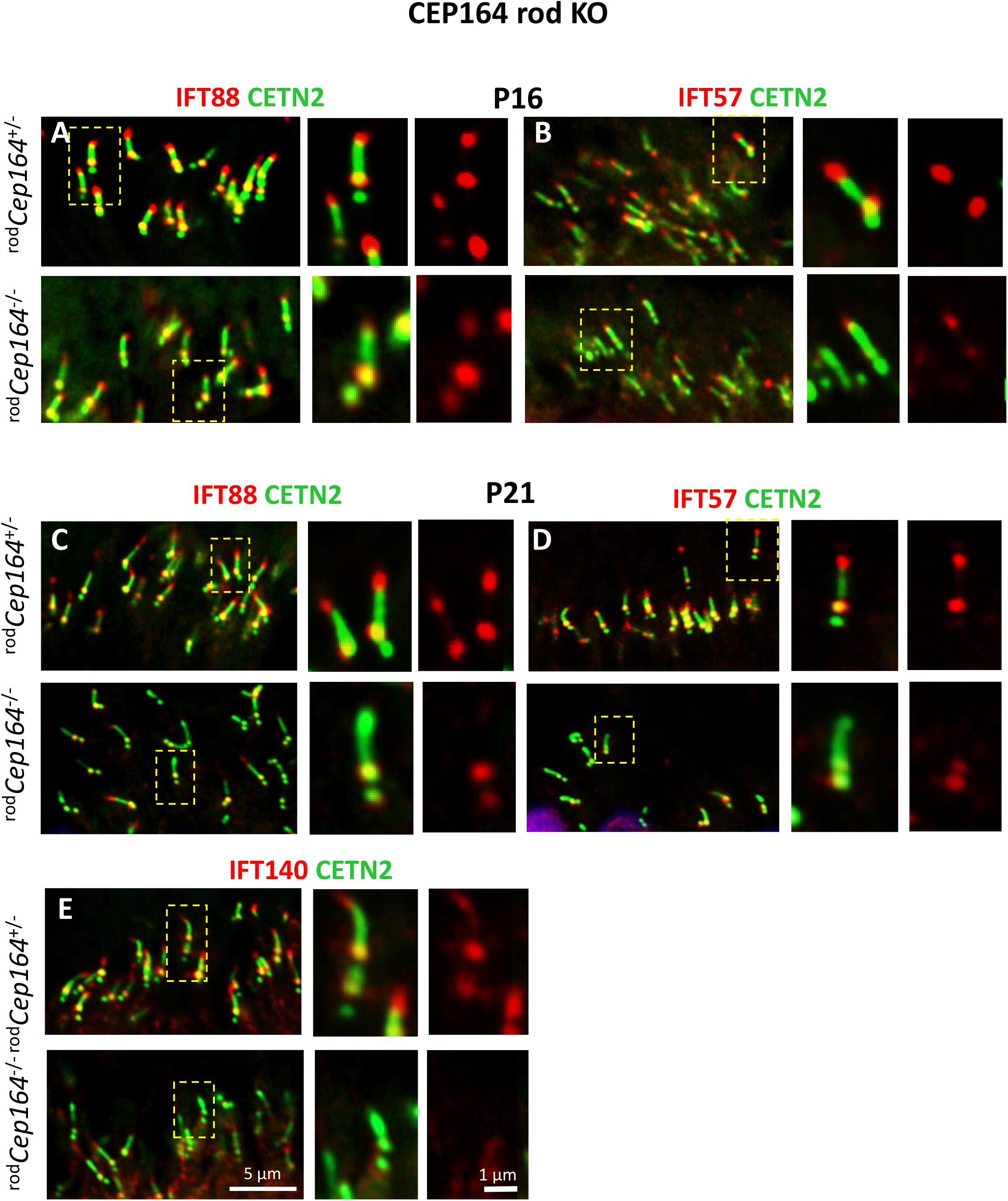
Loss of IFT88, IFT57 and IFT140 in ^rod^*Cep164*^-/-^ photoreceptors. **A, B**, immunohistochemistry of ^rod^*Cep164*^+/-^ (upper panels) and ^rod^*Cep164*^-/-^ (lower panels) cryosections (lower panels) at P16 probed with anti-IFT88 (A) and anti-IFT57 (B) antibodies. **C-E**, immunohistochemistry of P21 ^rod^*Cep164*^+/-^ (upper panels) and ^rod^*Cep164*^-/-^ (lower panels) cryosections (lower panels) probed with anti-IFT88 (C), anti-IFT57 (D), anti-IFT140 (E) and IFT122 (F) antibodies. Insets (right) show enlargements of individual representative BB/CC structures. Note decrease of IFT proteins at the CC tip and BB.

**Figure 9.**
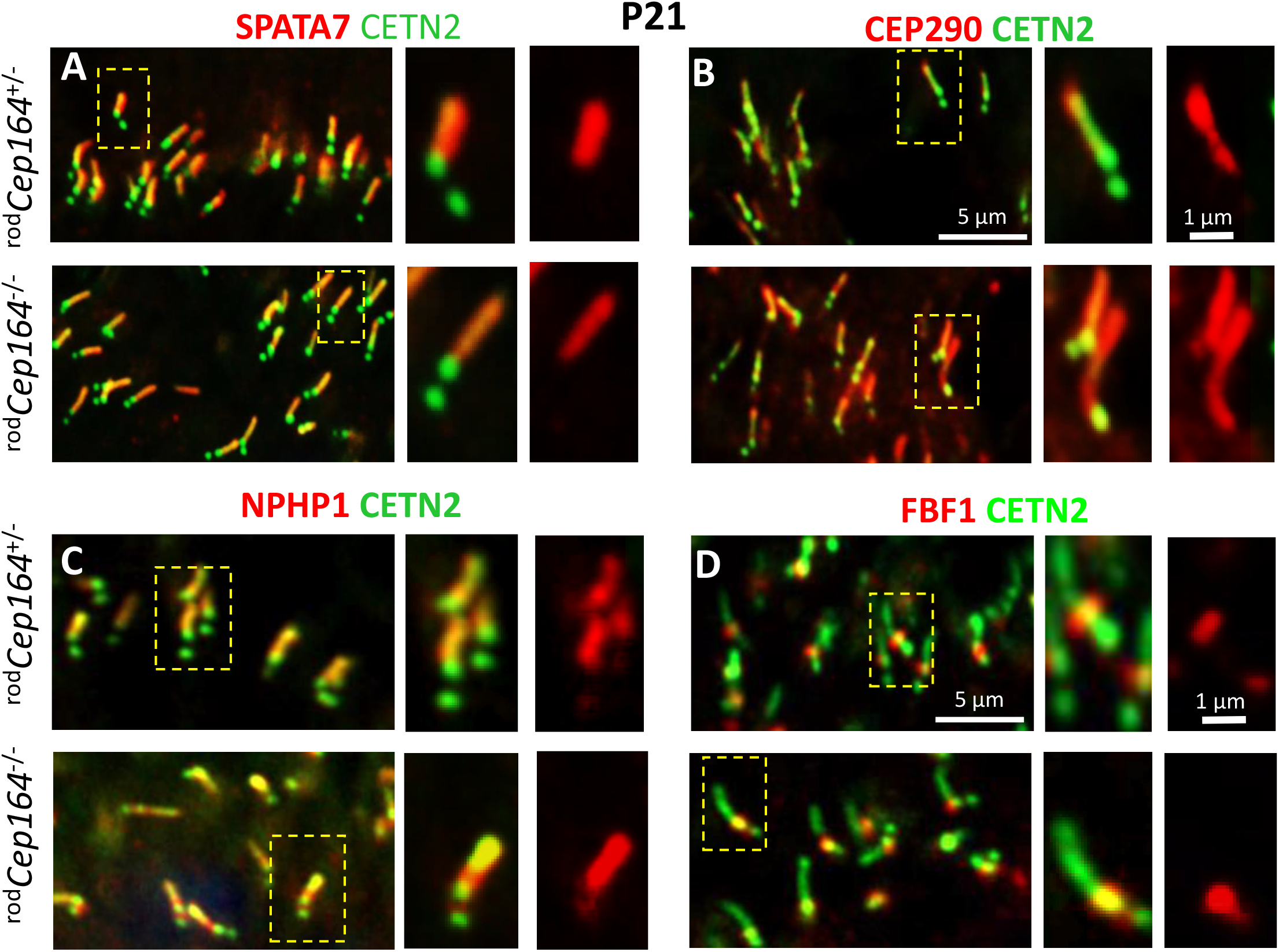
SPATA7, CEP290, NPHP1 and FBF1 are stable in the rod knockout. **A, B**, immunohistochemistry with anti SPATA7 and anti-CEP290. **C, D**, immunohistochemistry of ^rod^*Cep164*^+/-^(upper panels) and ^rod^*Cep164*^-/-^ cryosections (lower panels) at P21 probed with anti-NPHP1 (red) (C) and anti-FBF1 (D) antibodies. Note no relevant changes for SPATA7, CEP290, and NPHP1 localizations at the CC. FBF1 is a distal appendage protein locating at the BB. Insets (right) show enlargements of representative BB/CC structures identified by CETN2 or individual antibodies (red).

**Figure 10.**
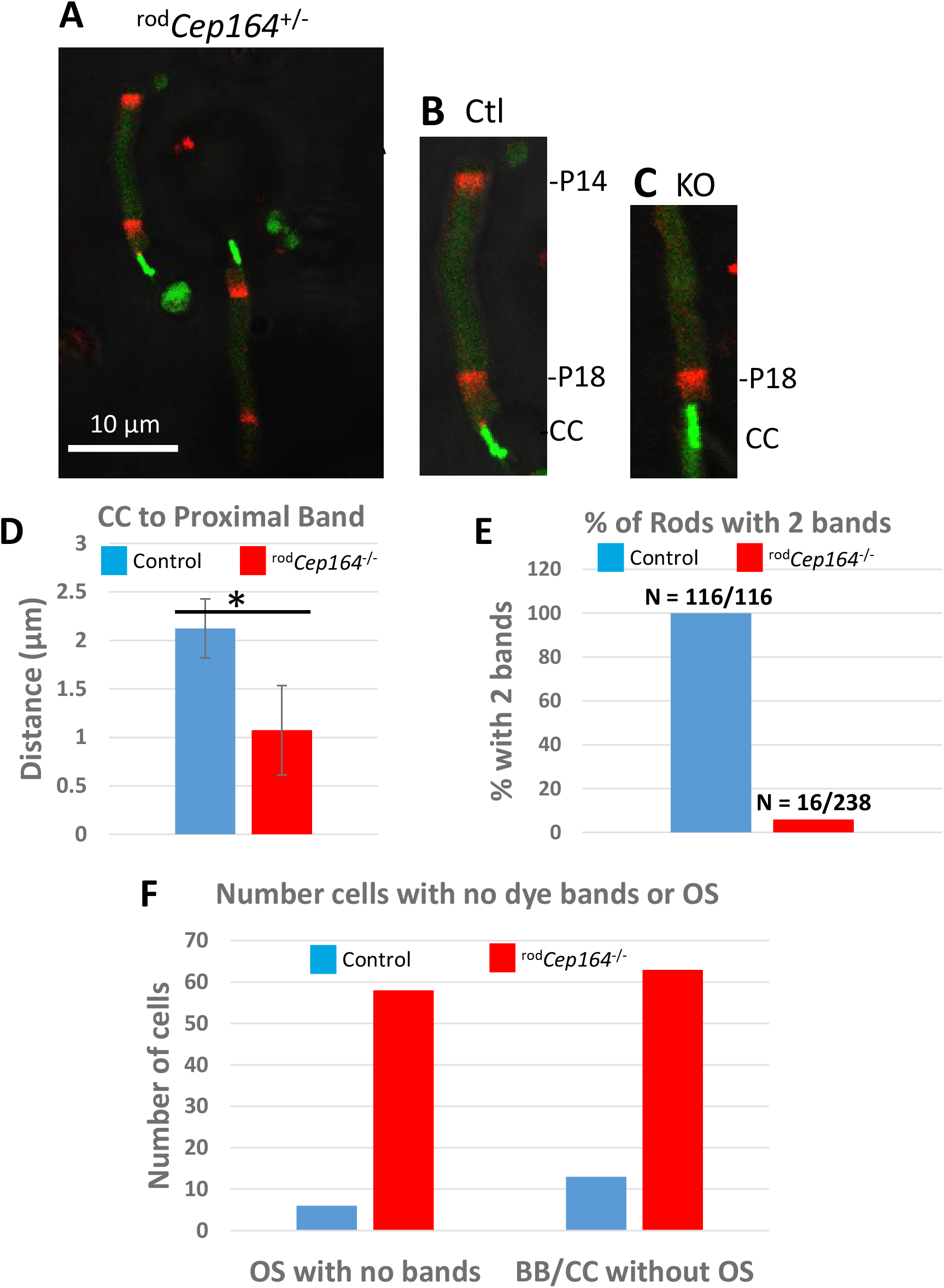
Intravitreal CF-568 dye injection. Dye injection of ^rod^*Cep164*^+/-^control versus ^rod^*Cep164*^-/-^ eyes on Egfp-Cetn2^+^ background reveals bands containing nascent discs. P19 retinas were collected, dissociated, and rods imaged. **A**, ^rod^*Cep164*^+/-^ control rods. **B**, single ^rod^*Cep164*^+/-^ rod with P14 and P18 disc bands. **C**, single ^rod^*Cep164*^-/-^ rod with P18 discs, but no P14 discs. **D**, statistical evaluation of the distance between CC-tip and center of P18 band in control (blue) and rod KO (red). **E**, % rods with P14 and P18 bands in control versus mutant. **F**, number of cells with no dye bands (left graph) and BB/CC structures without OS (right graph).

Distributions of two BB-associated proteins, pericentrin (PCNT) and ninein (NIN), were examined in ^rod^*Cep164*^-/-^ photoreceptors. PCNT interacts with numerous proteins including IFT proteins [37], the γ-tubulin ring complex [38], cytoplasmic dynein [39], PCM1 [40] and CEP215 [41], stabilizing microtubules and/or aiding microtubule nucleation [38]. PCNT locates to the pericentriolar material (PCM) surrounding the BB of mouse photoreceptors [38, 42]. Immunohistochemistry reveals that PCNT trails from the BB presumably along microtubules (**Fig. S4A**), upper panel); PCNT localization of P21 ^rod^*Cep164*^*-/-*^ BB is unchanged but slightly reduced (**Fig. S4A**, lower panel). Ninein colocalizes with γ-tubulin and C-Nap1 [43], and is important for positioning/anchoring the microtubule minus-ends to BB subdistal appendages [44]. Surprisingly, ninein locates only to the proximal axoneme of control photoreceptors (**Fig, S4B**, upper panels), whereas it labels robustly the proximal axoneme and BB/DC surround of ^rod^*Cep164*^*-/-*^ photoreceptors (**Fig S4B**, lower panels). Ninein was shown to be released from the centrosome and move along microtubules in epithelial cells [45]. The function of ninein at the proximal axoneme is unknown.

### Disc formation in ^rod^*Cep164*^*-/-*^ photoreceptors

We developed a fluorescent dye assay to monitor the formation of closed discs during disc morphogenesis. CF-568-hydrazide, a water-soluble, membrane impermeable, aldehyde-fixable red-fluorescent dye [46], was injected intravitreally at P14 (day 1) and P18 (day 4) and eyes harvested at P19 (day 5). Following retina dissociation, ^rod^*Cep164*^+/-^ control photoreceptors show two bands containing red fluorescent discs along the OS (**Fig. 10A, B**) indicating that CF-568 sequesters into the disc lumen during enclosure of a nascent disc. Nascent (P18) discs were located immediately adjacent to the CC (green, identified by EGFP-CETN2), whereas older (P14) discs are displaced distally towards the RPE, as shown by Richard Young [47, 48]. The two bands are separated by about 10 μm or ∼300 discs (**Fig. 10A**). Disc morphogenesis occurs as late as P18/19 in ^rod^*Cep164*^-/-^ OSs, albeit slower than controls as newly formed discs appear closer to the distal CC (**Fig. 10C, D**). Relative to control OS, P18 ^rod^*Cep164*^-/-^ bands are closer to the CC while P14 bands are very faint or absent, i.e., only ∼5% of KO rods display P14 bands (**Fig. 10E**). We identified more BB/CC without OS in the ^rod^*Cep164*^-/-^ versus control (∼5x more) (**Fig. 10F**), suggesting that a portion of P19 ^rod^*Cep164*^-/-^ OSs have already completely degenerated, and that loss of OSs is not primarily due to mechanical dissociation.

### CEP164 deletion by tamoxifen induction

To further explore consequences of CEP164 depletion, we deleted *Cep164* in *Cep164*^F/F^;Prom1-CreER^T2^ one-month old mice by tamoxifen induction. Prom1 expression in adult mice occurs in rod/cone photoreceptors along with cells of the brain, pancreas, intestine/colon, kidney, lung and reproductive system (male and female) [25]. *Cep164*^F/+^;Prom1-CreER^T2^ mice (**Fig. 11A**, left columns), *Cep164*^+/+^;Prom1-CreER^T2^ mice (**Fig. 11**, middle columns) and *Cep164*^F/F^;Prom1-CreER^T2^ mice (^tam^*Cep164*^-/-^) (**Fig. 11**, right columns) were injected with tamoxifen at ages ranging from P28 to P30. Intraperitoneal injection of tamoxifen for five consecutive days induced the nuclear translocation of *Cre*, and the degeneration rate was assessed in retina cryosections of eyes harvested 2-4wPTI after the first injection.

**Figure 11.**
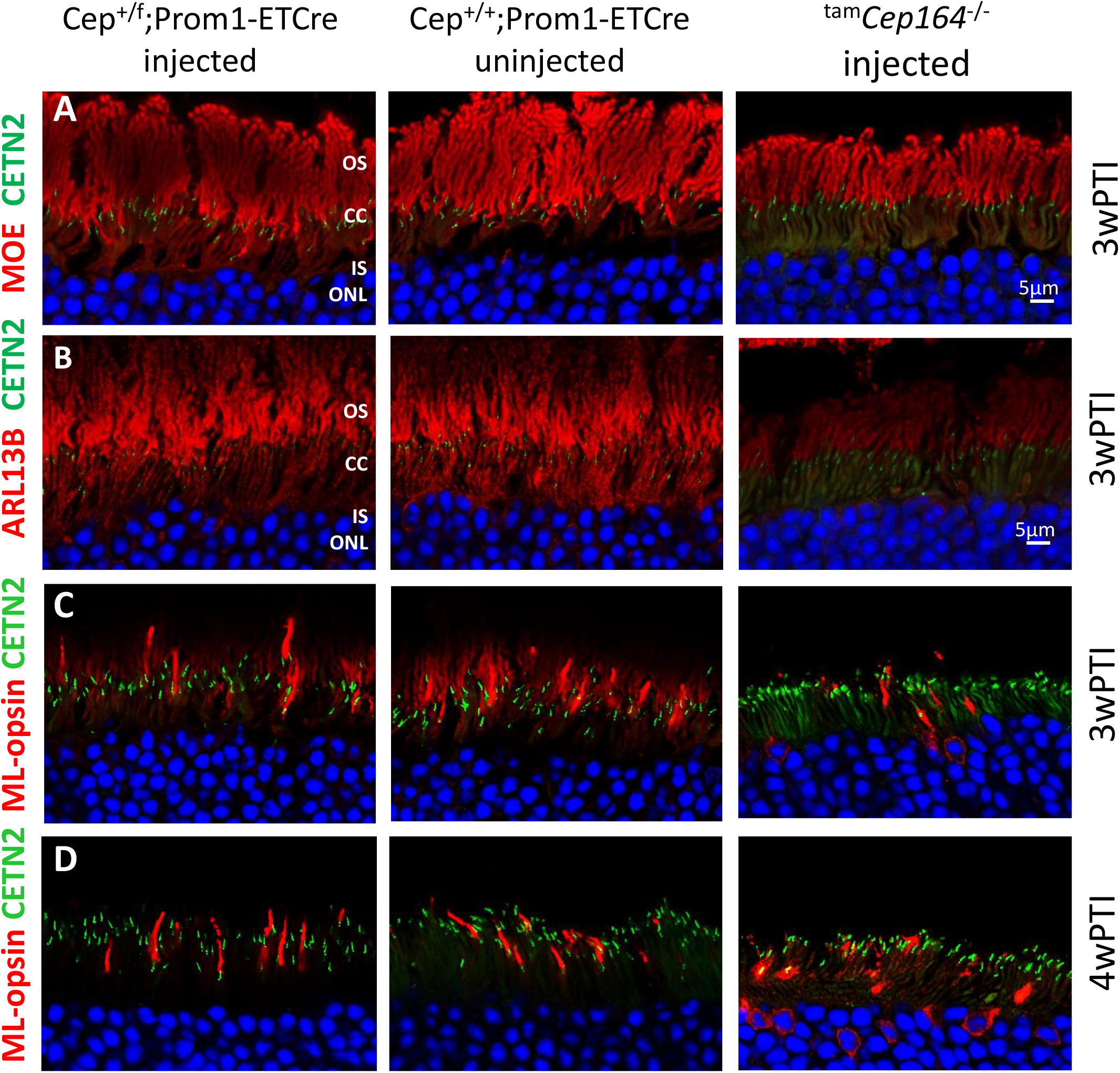
Tamoxifen-induced deletion of CEP164 in adult mouse. Rows **A-D**, Immunohistochemistry of cryosections of control *Cep164*^+/F^;Prom1-ETCre (injected) (left panels), *Cep164*^+/+^;Prom1-ETCre (uninjected) (middle panels) and *Cep164*^*F*/F^;Prom1-ETCre (^tam^*Cep164*^-/-^) (injected) (right panels) probed with MOE (anti-PDE6) (A), with anti-ARL13B (B), with ML-opsin antibody (C, D). Rows A-C are at 3wPTI, row D at 4wPTI.

Labeled by anti-PDE6 antibody (MOE), ^tam^Cep164^-/-^ rod OSs degenerated slowly and were approximately half-length at 3wPI (**Fig. 11A**, right panel) relative to control OS (left and middle panels). ^tam^Cep164^-/-^ rod OSs probed with anti-ARL13B (**Fig. 11B**, right panel) revealed that ARL13B is unable to reach the OS in the absence of CEP164, confirming the interaction of these two proteins [22]. ^tam^Cep164^-/-^ cone OS identified by anti-ML-opsin were shorter at 3wPTI (**Fig, 11C**, right panel) compared to controls (**Fig. 11C**, left and middle panels) and ML-opsin mislocalized in part to the cone IS. At 4wPTI, mutant cone OSs were largely degenerated (**Fig. 11D**, right panel). Of 340 ^tam^Cep164^-/-^ rods analyzed by confocal microscopy at 2.5wPTI, one-half had CEP164 associated with the BB, while the other half was CEP164-free (**Fig. 12A**) indicating a 50% induced knockout efficiency. Anterograde IFT57 and IFT88 showed normal distributions at control BB and CC tips (**Fig. 12B, C**, upper panels) as observed in the CEP164 rod knockout at P16 (**Fig. 8A, B**). As expected, IFT57 and IFT88 were absent at the ^tam^*Cep164*^-/-^ CC tip and reduced at BB. NPHP1 localization at CC was unaffected (**Fig. 12D**).

**Figure 12.**
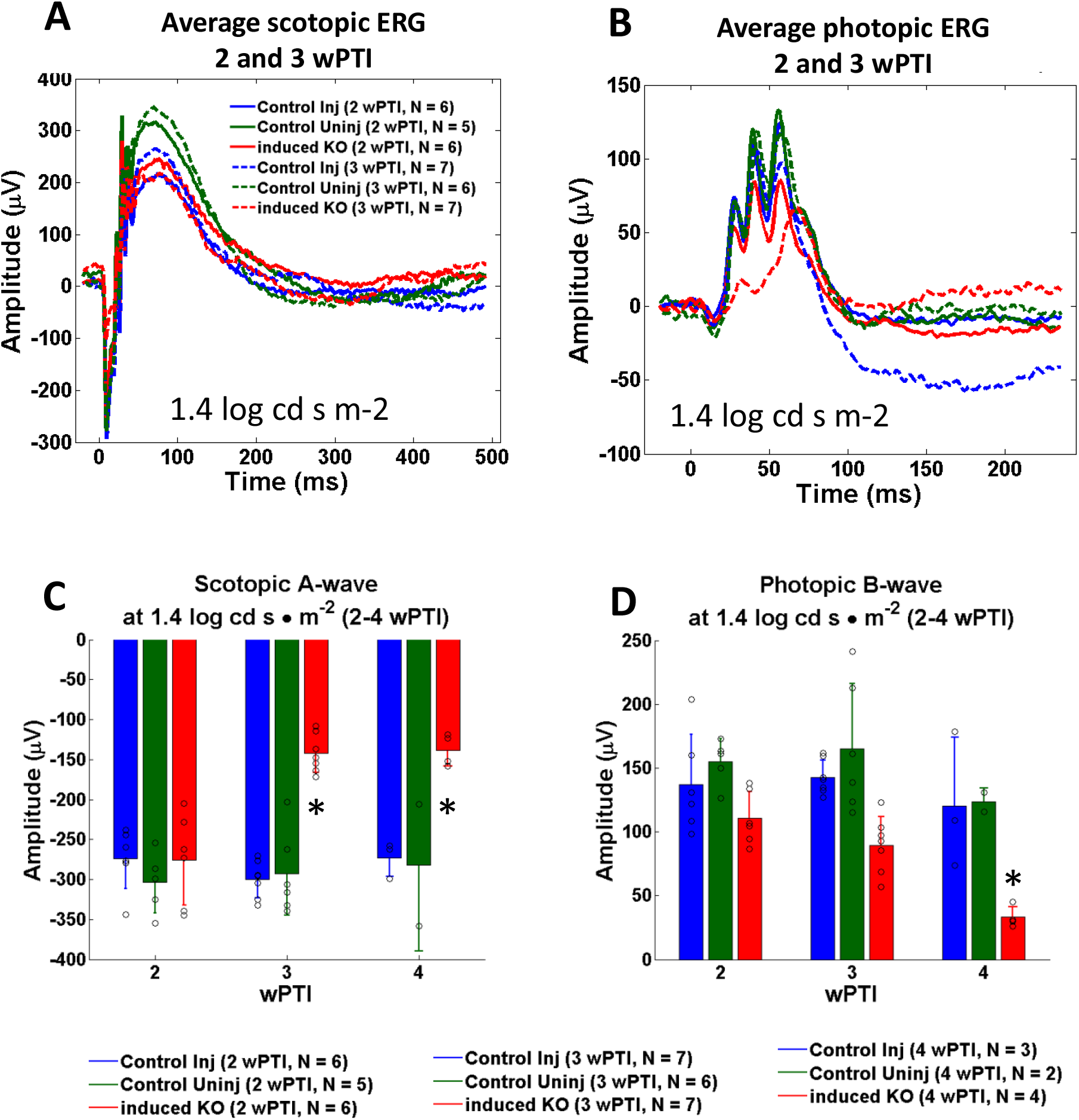
Immunohistochemistry with anti IFT57, IFT88 and anti-NPHP1 in ^tam^*Cep164*^-/-^ mice. **A**, BB of injected controls (no Cre), injected hets (with Cre), uninjected control and ^tam^*Cep164*^-/-^ retina assayed for presence of CEP164 immunoreactivity. Essentially all control BB displayed CEP164; in ^tam^*Cep164*^-/-^ retina, 170 of 340 BB did not. **B-D**, immunohistochemistry of 2.5wPTI ^tam^*Cep164*^+/-^ (upper panels) and ^tam^*Cep164*^-/-^ (lower panels) photoreceptors probed with anti-IFT57 (red) (B), anti-IFT88 (C) and anti-NPHP1 (D) antibodies. IFT57 and IFT88 immunolabeling are reduced in the knockout. Insets (right) show enlargements of representative BB/CC structure identified by EGFP-CETN2 (green) or individual antibodies (red).

Average ^tam^*Cep164*^*-/-*^ scotopic a-wave amplitudes, not significantly reduced at 2wPTI, were diminished by 3wPTI (**Fig. 13A**). Statistical evaluation of scotopic a-waves at 2-4wPTI at 1.4 log cd s/m^2^ confirmed diminished a-wave amplitudes (50% and 45%, respectively) (**Fig. 13C**). A-wave amplitudes did not significantly change at a higher light intensity of 2.4 log cd s/m^2^ (**Fig. S5A**). By contrast, ^tam^*Cep164*^*-/-*^ photopic b-waves persistently diminished at 2-4wPTI down to 40% of control at 4wPTI (**Fig. 13B, D**). A- and b-wave amplitudes were not reduced further even at 6wPTI and 9wPTI (**Fig. S6**). Pan-retina ERG as a function of light intensity (−4.5 to 2.4 log cd s/m2) did not reveal significant differences of scotopic/photopic a- and b-wave amplitudes at 2wPTI (**Fig. 13**). At 3wPTI, scotopic ERG showed diminished a- and b-wave amplitudes beginning at 0.6 log cd s/m2 (**Fig. S7**). Photopic b-wave amplitudes of control and ^tam^*Cep164*^*-/-*^ mice were statistically identical at 2wPTI, but decreased by 3wPTI.

**Figure 13.**
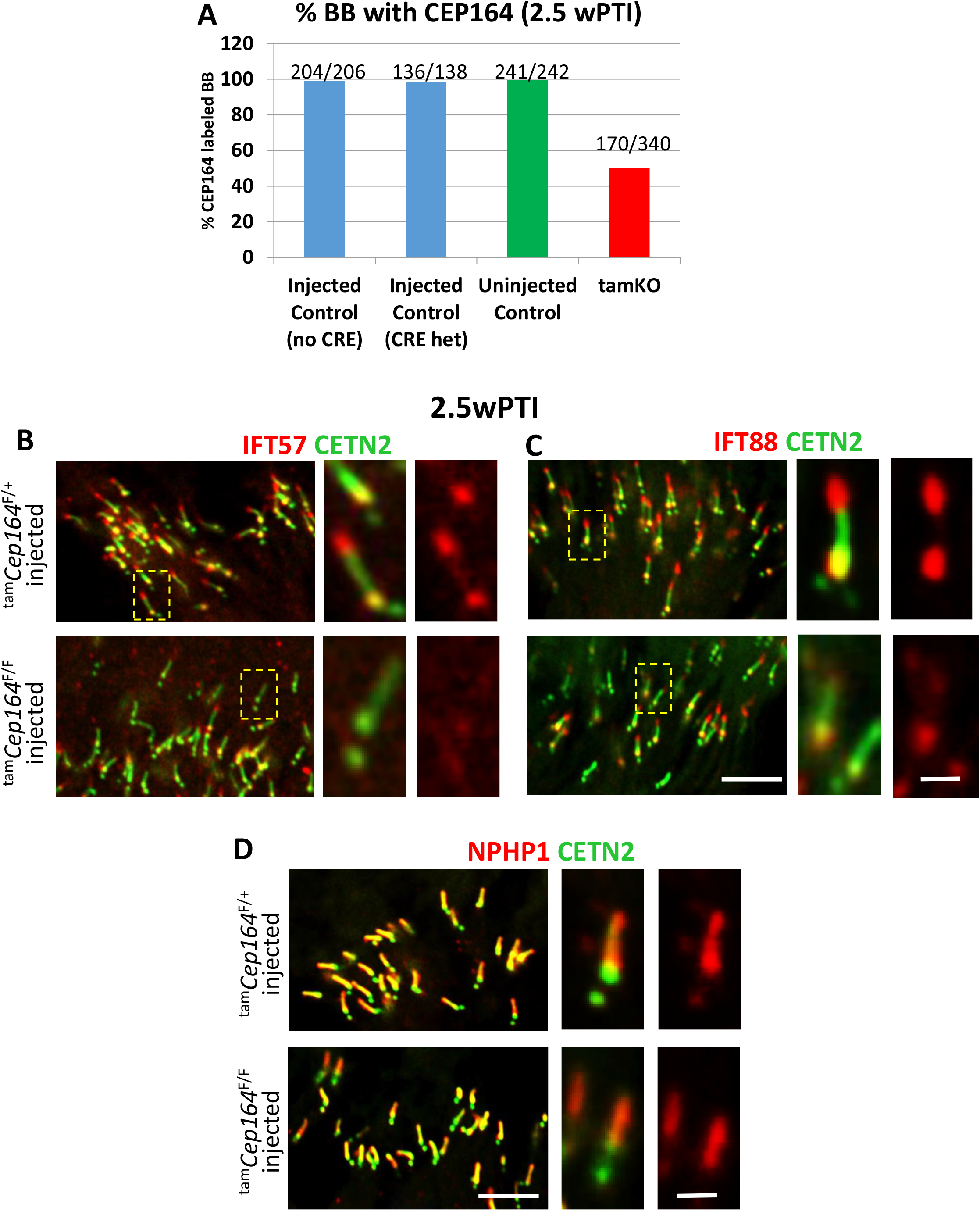
Electroretinography of control and ^tam^*Cep164*^-/-^ mice. **A, B**, average scotopic (A) and photopic ERG (B) at 1.4 log cd s m^-2^. 2wPTI is shown in solid traces, and 3wPTI in hatched traces. Control-injected traces (blue), control-uninjected traces (green), ^tam^*Cep164*^-/-^ (red). Traces are averaged with N = 5-7. **C, D**, statistical evaluation of scotopic a-waves (C) and photopic b-wave amplitudes (D) at 2wPTI, 3wPTI and 4wPTI at 1.4 log cd s/m^2^. Colors as in A and B, number of animals (N) used is shown at the bottom.

## Discussion

Many cell culture studies address the role of CEP164 during BB docking and early ciliogenesis [2, 12, 30, 41] where disruption of CEP164 blocked primary cilium formation [12, 49]. Similar results were obtained in zebrafish [12, 13, 16] and mouse [1, 17, 50]. Here, we describe effects of CEP164 deletion at three different stages of retina development using conditional recombination using Cre drivers which express Cre beginning at E9 (Six3Cre) [23], in the second postnatal week (iCre75) [24] or in the adult mouse (tamoxifen injection). Two drivers express Cre after ciliogenesis (iCre75 and tamoxifen inducible Prom1-CreER^T2^) when CEP164 is bound to the distal appendages and the BB is docked to the apical membrane.

Six3Cre-based deletion of CEP164 has no effect on pre-ciliogenesis photoreceptor development, ONL lamination (**Fig. 1H**) or retina morphology (**Fig. 2B**), but prevents BB docking (**Fig. 2D**) and OS formation (**Fig. 3B, E**), compatible with the well-documented role of CEP164 in cultured cells. Loss of CEP164 in developing retina leads to rapid rod and cone degeneration centrally with an LCA-like phenotype. Postnatal deletion of CEP164 with iCre75 is detectable after eye opening at P12, allowing the BB to mature and dock to the cell cortex. Control photoreceptor CC extend axonemes ∼P6, and OSs are mature at P21. Rod-specific CEP164 deletion begins when ciliogenesis and OS disc formation are active. In P15 and P21 ^rod^*Cep164*^*-/-*^photoreceptors (**Fig. 5B**), most BB have lost CEP164 but OSs are still congruent with BB-CC structure (**Fig. 5B, B’** and **Fig. 6B**). However, the P21 ^rod^*Cep164*^*-/-*^ ONL thickness is reduced and, at P30, only one nuclear row is present (**Fig. 6B**). PDE6 mislocalization was not obvious in the P21 ^rod^*Cep164*^*-/-*^ when OSs are disintegrating (**Fig. 6B**), suggesting delivery of PDE6 as OS discs continue assembly. The results suggest that the BB-CC-axoneme structure at the OS base is relatively stable even as the OS degenerates (**Fig. 6B**), in contrast to CEP164’s ciliogenesis role where CEP164 is required for recruitment and physical attachment of small preciliary vesicles to the mother centriole and membrane docking.

Loss of IFT88, IFT140, IFT57 and PCNT during deletion of CEP164 from P16-P21 (**Figs. 8 and S4**) implicates IFT malfunctioning as the cause of retina degeneration in ^rod^*Cep164*^*-/-*^ mice. Similar results were seen in tamoxifen-induction experiments at 2.5WPTI (**Fig. 12B, C**). IFT is a pathway integral for assembly, maintenance, and function of motile and primary cilia [51, 52]. IFT relies on molecular motors and IFT particles to move cargo along microtubules in anterograde (heterotrimeric kinesin 2) or retrograde (dynein-2) direction. IFT88 (Tg737) and IFT57 are required for ciliary and *Chlamydomonas* flagella assembly [53]. IFT57, discovered as Hippi involved in recruitment of caspase-8 in Huntington disease [54], stabilizes the IFT-B complex in *Chlamydomonas* flagella [55] and interacts with IFT20 and the kinesin-2 subunit, KIF3B [56]. Germline deletions of IFT88 are embryonically lethal [57, 58]. In mouse photoreceptors, a hypomorphic mutation in the IFT88 gene exhibited abnormal OS development and retinal degeneration demonstrating that IFT88 is important for OS maintenance [31]. In conditional retina IFT88 knockouts (^ret^*Ift88*^-/-^), BBs failed to extend CC and an axoneme [59].

Loss of IFT88 (Tg737) and IFT57 (Hippi) during CEP164 deletion confirms CEP164’s involvement in recruiting these proteins to the photoreceptor BB, first observed in primary cilia [30]. Another DAP protein involved in recruiting IFT-B proteins (IFT88 and IFT52) is C2CD3 (OFD14) which localizes to the distal ends of both mother and daughter centrioles. C2CD3 was shown to recruit IFT88 and IFT52 to the mother centriole [4]. As C2CD3 is required for the recruitment of DAP proteins including CEP164, the effect of recruiting IFT-B proteins may be indirect. Loss of C2CD3 results in shortened centrioles without appendages and failure of CP110 removal from the ciliary mother centriole, a critical step in initiating ciliogenesis.

Instability of the OS distal tip with continued disc morphogenesis proximally was also seen in the CF-568 dye assay that allows monitoring disc morphogenesis, enclosure and displacement in mouse rods (**Fig. 10**). Intravitreally-injected CF-568-hydrazide [46] is sealed into the disc lumen during the disc enclosure process. The number of labeled discs, limited by available dye, is fewer than 80, the number of discs generated per day. We propose that instability at the ^rod^*Cep164*^-/-^ OS distal tip may be caused by axonemal disintegration as IFT-B proteins dissociate from BB thereby impairing IFT (**Fig. 8**).

IFT cargo of rods is controversial. Current models suggest that rhodopsin and other integral membrane proteins traffic by anterograde IFT, mediated by kinesin-2, along the CC for deposition into nascent discs [60-63]. In our hands iCre75-based conditional knockout of KIF3A, a kinesin-2 obligatory subunit, did not prevent trafficking of rhodopsin and other proteins to the OS [64]. Conditional knockout of KIF3A with Six3Cre prevented CC and axoneme formation, although BB docked to the cell membrane correctly [59]. Conditional knockout of KIF3A and IFT88 in the adult mouse by tamoxifen injection, demonstrated that photoreceptor axonemes disintegrated slowly post-induction, starting distally, but rhodopsin and cone pigments trafficked normally for more than 2 weeks, a time interval during which the OS is completely renewed. Thus, visual pigments likely transport to the photoreceptor OS despite removal of KIF3A, IFT88 and impairment of IFT. Alternatively, kinesin-2-mediated anterograde IFT appears to be responsible for photoreceptor transition zone and axoneme formation [59].

CEP164 forms a complex with PDEδ (chaperone for prenylated protein), INPP5E (a farnesylated inositol polyphosphate 5 phosphatase) and ARL13B (GEF of ARL3) to traffic prenylated INPP5E to primary cilia in cell culture [22]. However, INPP5E of wildtype mouse photoreceptors localizes to the IS, is excluded from the OS, and is apparently independent of PDEδ [65]. Absence of INPP5E in the OS nullifies experiments to show that ARL13b, PDEδ or CEP164 act as OS chaperones for INPP5E. In ^ret^*Inpp5e*^-/-^ photoreceptors, CC and short rudimentary OS were established, but axonemal structure with discs failed to form [65]. ARL13B localizes to WT OS, but in ^ret^*Arl13b*^-/-^ [66] and ^ret^*Cep164*^-/-^ photoreceptors (**Fig. 3E**), CC and OS never form and ARL13B accumulates in the IS. In ^rod^*Cep164*^-/-^ photoreceptors, OSs form but ARL13B is barely detectable at P16 and absent at P21 (**Fig. 6D**), while in ^tam^*Cep164*^-/-^ photoreceptors, ARL13B is depleted in OS (**Fig. 11B**, right panel). These observations are consistent with [22] who show that ARL13B and CEP164 interact to enable ARL13B to reach the cilium, but not with [17] where ARL13B accumulated in CEP164-KO multiciliated cells. These results suggest that the functional network consisting of ARL13B, INPP5E, PDEδ and CEP164 is an arrangement of primary cilia for INPP5E and ARL13B to traffic to cilia, but is conserved only partially in photoreceptors.

## Summary

Deletion of CEP164 in photoreceptors before ciliogenesis prevented docking of BB to the IS cell membrane as shown in primary cilia. Deletion of CEP164 after ciliogenesis and BB docking to the IS cell cortex shows that the BB/CC/OS structure lacking CEP164 is initially stable. Post-ciliogenesis deletion of CEP164 causes loss of anterograde IFT proteins and impairment of IFT, an essential transport mechanism for assembly and OS maintenance. As OS proteins traffic to the ^rod^*Kif3a*^-/-^ OS [64] but the axoneme becomes unstable, we favor a model in which the ^rod^*Cep164*^-/-^ axoneme is not maintained as IFT-B proteins dissociate from the BB, causing OS degeneration as IFT trains fail. Loss of IFT88 and IFT57 during CEP164 deletion suggests that CEP164 recruits anterograde IFT proteins to the BB as proposed by [30], and identifies CEP164 and DAP structure as integral for initiating IFT.

## Author contributions

MR and WB designed the study; MR, GY, and JMF performed research; MR, GY, JMF, and WB analyzed data; KIT produced floxed mice; WB wrote the manuscript; MR, JMF, GY and KIT edited the manuscript.

## Acknowledgement

We thank Rui Du for help with dye injection, and Dr. Greg Pazour (University of Massachusetts Medical School) for providing IFT88, IFT157 and IFT140 antibodies. This work was supported by NIH grants R01HL139643 (KIT); EY08123, EY019298 (WB); EY014800-039003 (NEI core grant), 5T32 EY024234 (NEI training grant), by unrestricted grants to the University of Utah Department of Ophthalmology from Research to Prevent Blindness (RPB; New York) and by a grant from the Retina Research Foundation-Houston (Alice McPherson, MD). WB is recipient of the RPB Senior Investigator and Nelson Trust awards.

## Materials and Methods

### Animals

All procedures were approved by the University of Utah Institutional Animal Care and Use Committee (Protocol 18-11005). Prom1^tm1(cre/ERT2)Gilb^ (Stock No: 017743) and Egfp-*Cetn2* mice Stock No. 008234 - CB6-Tg(CAG-EGFP/CETN2)3-4Jgg/J) were obtained from The Jackson Laboratory. iCre75 mice were generated in Utah [24]. Cep164F/F mice were described previously [17].

### Generation of retina- and rod-knockout mice

*Cep164*^F/F^ mice were crossed with Six3Cre [23] or iCre75 transgenic mice [24] kept on Egfp-Cetn2 background to generate *Cep164* ^F/+^;Six3Cre;Egfp-*Cetn2* (^ret^*Cep164*^+/-^) or *Cep164* ^F/+^;iCre75;Egfp-*Cetn2* (^rod^*Cep164*^+/-^) mice. Mice were then backcrossed to *Cep164*^F/F^ to generate experimental animals.

### Generation of knockouts by tamoxifen induction

*Cep164*^F/F^ mice were bred to Prom1-CreER^T2^ mice [25] to generate *Cep164*^F/+^;Prom1-CreER^T2^ mice. Expression of CreER^T2^ is driven by the prominin 1 (Prom1) promoter/enhancer. *Cep164*^F/+^;Prom1-CreER^T2^ mice were bred to *Cep164*^F/F^ mice to generate mice for tamoxifen injection. Tamoxifen was administered via intraperitoneal injection in adult mice (P28-P30 at time of first injection). Tamoxifen was dissolved in corn oil to a stock solution of 20mg/ml. Mice were dosed with 150 mg/kg body weight for 5 consecutive days according to their weight on the first day of injections (i.e., 7.5 µL of 20mg/ml tamoxifen solution per gram weight). ERG and eye collection for confocal immunolocalization were performed 2-9 weeks after the first injection.

### Genotyping

Genomic DNA was extracted from fresh tissue by dissolving tail clips or retina DNA from P8-12 day old mice in 150 µL tail lysis buffer at 50-60°C for 1-2 hour. Digests were centrifuged at 15000 rpm for 5 minutes. Supernatant was added to an equal volume of isopropanol and centrifuged at 15000 rpm for 5 minutes. The DNA pellet was rehydrated in 75 µL H_2_O. Genotyping was achieved by polymerase chain reaction with EconoTaq® DNA polymerase (Lucigen). Primers for genotyping of CEP164 mice: 5’-CCATCTGTCCAGTACCATTAAAAA and 5’-CCCAGAATACAACATGGGAGA (WT allele, 215 bp; floxed allele, 415 bp). The CEP164 exon 4 excision assay used CEP164 floxed allele F1 primer, 5’- CCATCTGTCCAGTACCATTAAAAA and R2 primer 5’-GACAAGTTCCATCTACCACAATC (WT allele 1525 bp; Floxed allele 1696 bp; Exon 4 excision allele 719 bp).

### Dye injection

CF-568-hydrazide, a water soluble, membrane impermeable, aldehyde-fixable red-fluorescent dye [46]. Heterozygous control (^rod^Cep164^F/+^;Egfp-Cetn2) and knockout (^rod^Cep164^F/F^;Egfp-Cetn2) eyes were injected intravitreally with 1.5 µL of 0.5% CF-568-hydrazide dye [46] in 1x PBS on P14 and P18, and retinas were harvested on P19, at which time about half OS-length has been replaced. Treated retinas were dissociated mechanically and images acquired using a Zeiss LSM 800 microscope.

### Immunohistochemistry

Eyes were either embedded without fixation, or fixed 10 min or 1 hour in 4% paraformaldehyde in 0.1M phosphate buffer (pH 7.4), and embedded in OCT compound (Fisher). Sections were allowed to adhere to slides for 30 minutes at 37°C. Eyes that had no initial fixation had their OCT compound dissolved by adding a small amount of 1X PBS and holding stationary for 5 min at room temperature. Then 1:1 methanol-acetone was added for 10 min at 4°C. Slides were rehydrated by washing either 10 min (X3) or 5 min (X4) in 1X PBS. All sections used were from the central retina near the optic nerve, except when labeled as peripheral. Sections were blocked in 5% normal goat serum (NGS); 0.1% Triton in 1X PBS (for 1 hour 4% PFA fixation) or 5% NGS in 1X PBS (for 10 min 4% PFA or 10 min methanol-acetone fixation) for 1 hour. Antibodies, fixation, dilutions and sources: PDE6 (1 hour 4% PFA, 1:1000; MOE Cytosignal) [64]; OPN1MW/OPN1LW (1 hour 4% PFA, 1:500, Millipore Sigma AB5405) [65]; CEP164 (10 min 4% PFA, 1:350, Sigma-Aldrich); ARL13B (1 hour 4% PFA,1:350, ProteinTech); IFT57 (10 min 4% PFA, 1:200, Pazour lab); IFT88 (10 min 4% PFA, 1:200, Pazour lab); IFT120 (10 min 4% PFA, 1:150, Pazour lab); SPATA7 (10 min 4% PFA, 1:100, Abcam, purified as described [34]); CEP290 (10 min 4% PFA, 1:300, Swaroop laboratory); PCNT (10 min 4% PFA, 1:250, Covance); Ninein (10 min 4% PFA, 1:100, Abclonal); NPHP1 (10 min 4% PFA, 1:100, [67]); FBF1 (10 min methanol-acetone fix, 1:200, Proteintech). Primary antibodies were diluted in blocking buffer and applied to sections; sections were then incubated overnight at 4°C. Slides were washed for 10 min (X3) or 5 min (X4) in 1X PBS. Secondary antibodies were diluted in blocking buffer (goat anti-rabbit Alexa Fluor 555, 1:1000 (Invitrogen 32732); goat anti-mouse Alexa Fluor 555, 1:1000 (Invitrogen 32737); DAPI, 1:3000), applied to the sections and incubated in motion, in the dark, for 1 hour at room temperature. Slides were washed for 10 min (X3) or 5 min (X4) in 1X PBS. Slides were dipped briefly in deionized H_2_O, and coverslipped using Fluoromount-G® Mounting Medium (Southern Biotech). Images were acquired using a Zeiss LSM 800 confocal microscope with 63X objective and post-processed with Airyscan. All genotypes of a given age and antibody were imaged at a single z-plane using identical settings for laser intensity and master gain. Digital gain was 1 for all images. Pinhole size was set for 1AU on the red channel (39 µm for the 40X objective). Post-processing of non-saturated images consisted of equal adjustments to brightness/contrast of control and knockout images using Adobe Photoshop but without affecting the conclusions made. Red channel separation was obtained by isolating the R-channel in the “blender options” of Adobe Photoshop.

### Electroretinography

Scotopic and photopic ERG measurements were performed using P16 and P25 for Six3Cre experiments and P16, P21, or P30 mice for the iCre75 experiments and at 2, 3, 4, 6 or 9 weeks post-tamoxifen induction for the Prom1-CreER^T2^ experiments. Prior to ERG the mice were dark-adapted overnight and anesthetized with intraperitoneal injection of 1% ketamine/0.1% xylazine at 10 µl/g body weight. The mice were kept warm during ERG by using a temperature-controlled stage. Scotopic and photopic responses were recorded as described [66] using a UTAS BigShot Ganzfeld system (LKC Technologies, Gaithersburg, MD). Scotopic single-flash responses were recorded at stimulus intensities of -4.5 log cd s·m^-2^ [log candela seconds per square meter] to 2.4 log cd s·m^-2^). Mice were light-adapted under a background light of 1.48 log cd s·m^-2^ for 5min prior to measuring photopic responses. Photopic single-flash responses of control and knockout were recorded at stimulus intensities of -0.1 log cd s·m^-2^ to 1.9 log cd s·m^-2^.

### Statistical analysis

We performed an unbalanced two-factor ANOVA to compare experimental and control animals for their quantified A- and B-wave ERG response across multiple ages or weeks post-tamoxifen induction. Post-hoc multiple comparison was performed using Tukey’s honestly significant difference criterion. Statistical significance was determined using an alpha value of p < 0.05. All ERG statistics were computed using MATLAB’s statistical toolbox “anovan” and “multcompare” functions.

We performed a two-tailed Student’s *t*-Test assuming equal variances to compare ONL thickness for ^ret^*Cep164*^+/-, ret^*Cep164*^-/-, rod^*Cep164*^+/-^ and ^rod^*Cep164*^-/-^. Retina measurements used for these calculations were determined based on an average of three measurements per retina. Microsoft Excel function “T.TEST assuming equal variances, two-tailed” was used to calculate the p-value, with statistical significance determined using an alpha value of p < 0.05.

## Supporting information

### Supplemental Figures

**Figure S1. Localization of CEP164 within BB docked to inner segment membrane (ciliogenesis). A**, immunolocalization of CEP164 (red) in P21 *Cep164*^+/+^;Egfp-Cetn^2^ (green) photoreceptors. CEP164 is an inner segment protein associating with BBs. **A’**, BB enlargement with CC and daughter centriole (DC). Expression of transgenic EGFP-CETN2 identifies the CC, BB (= mother centriole, MC) and DC. **B**, BB docking pathway. **a**, centrosome consisting of MC and DC; MC and DC distal ends are capped by CP110/CEP97 (green). **b**, MC with distal appendage proteins (DAPs, light blue) and CEP164 (red). **c**, TTBK2 (*tau* tubulin kinase 2) phosphorylates DAPs which removes the terminal cap and allows docking. MC matures to BB. **d**, BB cap removal permits extension of A- and B-tubules to form axoneme and the CC.

**Figure S2. Generation of *Cep164* conditional knockout mice. A**, schematic of human centrosomal protein 164 kDa (CEP164, NPHP15) (upper panel) and mouse CEP164 (lower panel) consisting of a WW domain (yellow) and three coiled-coiled domains (CC). Positions of human mutations associated with nephronophthisis (NPHP)/retina degeneration, and truncation point (asterisk) of mouse CEP164 in the retina knockout are indicated. **B**, schematic of *Cep164 ex*ons 1-7 with loxP recombination sites flanking exon 4. The FRT site originates from removing the gene trap with flippase [17]. **C**, Cre-induced recombination (achieved with Six3Cre, iCre75, or Prom1-ETCre) results in excision of exon 4 and generation of a null allele.

**Figure S3. Genotyping assays. A-E**, PCR genotyping for the presence of WT and floxed Cep164 alleles (A), as well as Six3Cre (B), iCre75 (C), CETN2-EGFP (D), and Prom ETCre (E) transgenes from tail DNA. **F**, verification of exon 4 excision using primers, F1 and R2, and P6 ^ret^*Cep164*^*-/-*^ retina DNA as template. **G**, immunohistochemistry of P6 WT (upper panel) and ^rod^*Cep164* ^-/-^ cryosections (lower panel) with anti-CEP164 (red). Note centrioles but absent CC in the knockout. **H**, rod knockout verification of exon 4 excision using primers, F1 and R2, and P16 ^ret^*Cep164*^*-/-*^ retina DNA as template. Lane 1, size markers; lane 2, +/F DNA; lane 3, F/F DNA; lane 3, +/F DNA with iCre75; lane 5, F/F DNA and iCre75; lane 6, water control. Note knockout (-/-) is complete in lane 5. **I**, genotyping of tamoxifen-induced CEP164 deletion at 2.5 weeks post-tamoxifen injection (2.5wPTI).

**Figure S4. Immunohistochemistry with anti-PCNT and anti-NIN. A, B**, immunohistochemistry of P21 ^rod^*Cep164*^+/-^ (upper panels) and ^rod^*Cep164*^--/-^ cryosections (lower panels) probed with antibodies (red) directed against PCNT (A) and NINEIN (B). Pericentrin locates to the pericentriolar material surrounding the BB (yellow); PCNT levels appear slightly reduced in ^rod^*Cep164*^*-/-*^ (B). Ninein is a subdistal appendage protein weakly associated with centrioles and strongly with the proximal axoneme of control photoreceptors; centriolar (yellow) localization is much stronger in the rod knockout. Right, enlargements of representative BB/CC structures identified by CETN2 or individual antibodies (red).

**Figure S5. Evaluation of scotopic a-wave and photopic b-wave amplitudes of tamoxifen-induced knockouts versus controls. A, B**, statistical evaluation of scotopic a-wave (A) and photopic b-wave (B) amplitudes at 2, 3 and 4wPTI at 2.4 log cd s/m^2^ (scotopic) and 1.9 log cd s/m^2^. Control-injected traces (blue), control-uninjected traces (green), induced knockout (red). Number of animals (N) tested is shown, bottom.

**Figure S6. Statistical evaluation of scotopic a- and photopic b-wave amplitudes at 2-9wPTI. A-D**, scotopic a-waves (A, B) and photopic b-waves (C, D) at 1.4 lob cd s/m^2^ (A, C) and 2.4 log cd s/m^2^ (B, D) at 2-9 wPTI. That OS do not completely degenerate may be due to insufficient tamoxifen, or the PROM1 promoter unable to express higher levels of ETCre.

**Figure S7. Electroretinography as a function of light intensity. A-C**, amplitudes (μV) of scotopic a-wave (A), scotopic b-wave (B) and photopic b-wave (C) as a function of light intensity (log cd s m^-2^). Scotopic and photopic ERGs show statistically distinct responses at 3wPTI (^, p < .05).

**Figure S8. Model of photoreceptor CEP164-mediated transport. A**, During photoreceptor anterograde IFT, directed toward the microtubule plus-end, kinesin-2 (KIF3A, KIF3B and KAP subunits) moves cargo along the proximal axoneme (microtubule doublet). Cargo likely consists of IFT-A and -B particles, retrograde dynein 2 motors, axoneme building blocks and axoneme-stabilizing factors [59]. Rhodopsin [60, 61] and other OS integral membrane proteins may be additional cargo. Kinesin-2 may also move cargo to the axoneme tip along axonemal MT singlets, as KIF17 (Osm-3 in *C. elegans*) has no IFT function in photoreceptors [68]. Axoneme maintenance is currently unclear. Retrograde trafficking likely occurs by a minus-oriented dynein 2 motor that returns kinesin-2 and IFT particles to the BB. **B**, loss of IFT-A particles, caused by deletion of CEP164, shuts down IFT and photoreceptor OS degenerate.

